# Urbanization correlates with genetic and plastic variation of the spotted jewelweed flower morphology

**DOI:** 10.1101/2025.06.10.658901

**Authors:** Jérôme Burkiewicz, Julie Carvalho, Sophie Caporgno, Joëlle Lafond, Céline Devaux, Étienne Normandin, Simon Joly

## Abstract

The spectacular diversity of flowers is largely driven by pollinator-mediated selection that favors attractive flowers and effective pollen transfer. Urbanization has the potential to affect floral trait evolution by altering pollinator communities through environmental changes. Additionally, abiotic changes in urban habitats can induce phenotypic plasticity, further shaping evolutionary trajectories. We investigated how urbanization affects the genetic and plastic components of flower morphology of *Impatiens capensis* across four Canadian cities. We found that urbanization influenced the pollinator community composition and the body size of bumblebees, the species’ main pollinator, although the magnitude of the size effect varied among cities. Using a combination of field surveys and a common garden experiment, our results suggest that urbanization affects sepal size – a tubular floral organ in which pollinators enter to access the nectar – through both genetic and plastic responses. While plasticity sometimes masked the genetic determination of sepal size in the field, we observed a positive correlation between the genetic component of the sepal size and bumblebee body size. These results suggest that urban habitats may drive evolutionary changes in floral traits by modifying pollinator communities.

## Introduction

Flowers show an astounding diversity in shapes, sizes, colors, and scents across angiosperms, mostly resulting from selection imposed by pollinators (Galen, 1999; Harder & Johnson, 2009; Ollerton et al., 2011; Stebbins, 1970; van der Niet & Johnson, 2012; Whittall & Hodges, 2007). Flowers can genetically adapt to changes in pollinator communities or abundance by evolving traits that enhance attraction, such as color, scent, and nectar, that facilitate pollen transfer by improving the mechanical fit with pollinators through shape or size modifications, or that promote self-pollination (Acoca-Pidolle et al., 2024; Gervasi & Schiestl, 2017; Gómez et al., 2009). But flowers can also be modified by their environment through phenotypic plasticity (Caruso, 2006; Gómez et al., 2020), which can be either adaptive or maladaptive depending on whether the environment modifies the phenotype in a way that increases or decreases the mean fitness of the population, respectively (Dorey & Schiestl, 2022). Also, phenotypic plasticity can hide genetic adaptation if it affects the phenotype in a way that opposes natural selection, a concept called counter-gradient variation (Conover et al., 2009; Conover & Schultz, 1995). Despite the important roles of genetics and phenotypic plasticity in determining phenotypes, few studies have tried to tease them apart when investigating evolutionary responses of flower morphology across environmental gradients.

Urban habitats offer a unique opportunity to study evolutionary responses to environmental changes (Johnson & Munshi-South, 2017). The development and spread of cities alter abiotic conditions by remodeling the physical environment and biotic conditions through their effects on species richness, abundance, interactions, and evolution (Alberti et al., 2017; Johnson & Munshi-South, 2017; McKinney, 2008; Santangelo et al., 2022). Furthermore, since urban habitats are generally more similar to each other than to their surrounding natural environments (Santangelo et al., 2022), studying species response to urbanization in different cities is equivalent to a replicated evolutionary experiment (Santangelo et al., 2020), and could uncover parallel or idiosyncratic responses across cities (Diamond et al., 2018; Santangelo et al., 2022).

Urbanization is expected to disrupt mutualistic interactions, such as pollination, because these interactions require compatibility between flowering and pollinator activity and phenotype matching between the plant and pollinator traits (Fisogni et al., 2020; Irwin et al., 2020). Pollinator communities commonly exhibit changes along urbanization gradients (Baldock et al., 2019; Hall et al., 2017; Theodorou et al., 2020; Wenzel et al., 2020), but urban habitats could also affect the body size of insect pollinators via changes in the pollinator species composition or through evolutionary responses to changes in resource availability and temperature (Goulson, 2010). So far, studies have produced conflicting results regarding how urbanization affects bumblebee sizes (Eggenberger et al., 2019; Theodorou et al., 2021). A greater distance between resource patches, as commonly found in cities, could favor larger insects with greater flight capacities (Greenleaf et al., 2007). On the other hand, high temperatures could favor smaller insects with greater thermoregulation potential (Eggenberger et al., 2019). Such changes in pollinator body size, as well as those in the abiotic environments, can potentially affect flower morphology through both genetic and plastic responses. Yet, few studies have investigated the impact of urbanization on the evolution of flower traits while also considering changes in pollinator communities (Irwin et al., 2018; Rivkin et al., 2020; Ushimaru et al., 2014) and attempted to disentangle genetic from plastic responses (but see Irwin et al., 2014).

We tested whether urbanization impacts, directly or indirectly through its effects on pollinator communities, the size and shape of flowers of the spotted jewelweed, *Impatiens capensis* Meerb. This annual species has a mixed pollination strategy, bearing both closed, green, and tiny self-fertilizing (cleistogamous) flowers, and, open, orange, large protandrous and mainly outcrossing (chasmogamous) flowers (Fig. 1). This reproductive strategy is plastic, as a greater proportion of chasmogamous flowers are produced with more light (Schemske, 1978; Waller, 1980), but it does not seem to be affected by urbanization (Barker & Sargent, 2020). Populations of *I. capensis* growing in sunlight produce numerous chasmogamous flowers pollinated by bumblebees, other bees, wasps, flies, and hummingbirds (Fenster et al., 2004; Rust, 1977, 1979), which all feed on pollen or the nectar secreted in the spur. The spur curvature of *I. capensis* is heritable, and affects pollen removal by hummingbirds (Travers et al., 2003) and the duration of visits by bees (Young, 2008). *Impatiens capensis* evolved smaller sepals under relaxed pollinator-mediated selection (Zhao & Schoen, 2022), and larger flowers under pollinator limitation (Panique & Caruso, 2020). Also, natural populations in Montreal, Canada, had larger sepals and longer nectar spurs compared to their urban counterparts (Faure et al., 2023). Finally, reproductive success was shown to be limited by pollen deposition in urban and natural habitats (Barker & Sargent, 2020), potentially allowing natural selection to drive floral phenotypic changes (Harder & Aizen, 2010). Therefore, the generalist pollination strategy of *I. capensis*, and its reported capacity to respond to its pollinator communities in both urban and natural habitats make it an excellent system for studying the evolution of flower shape and size in response to urbanization.

**Fig. 1.**
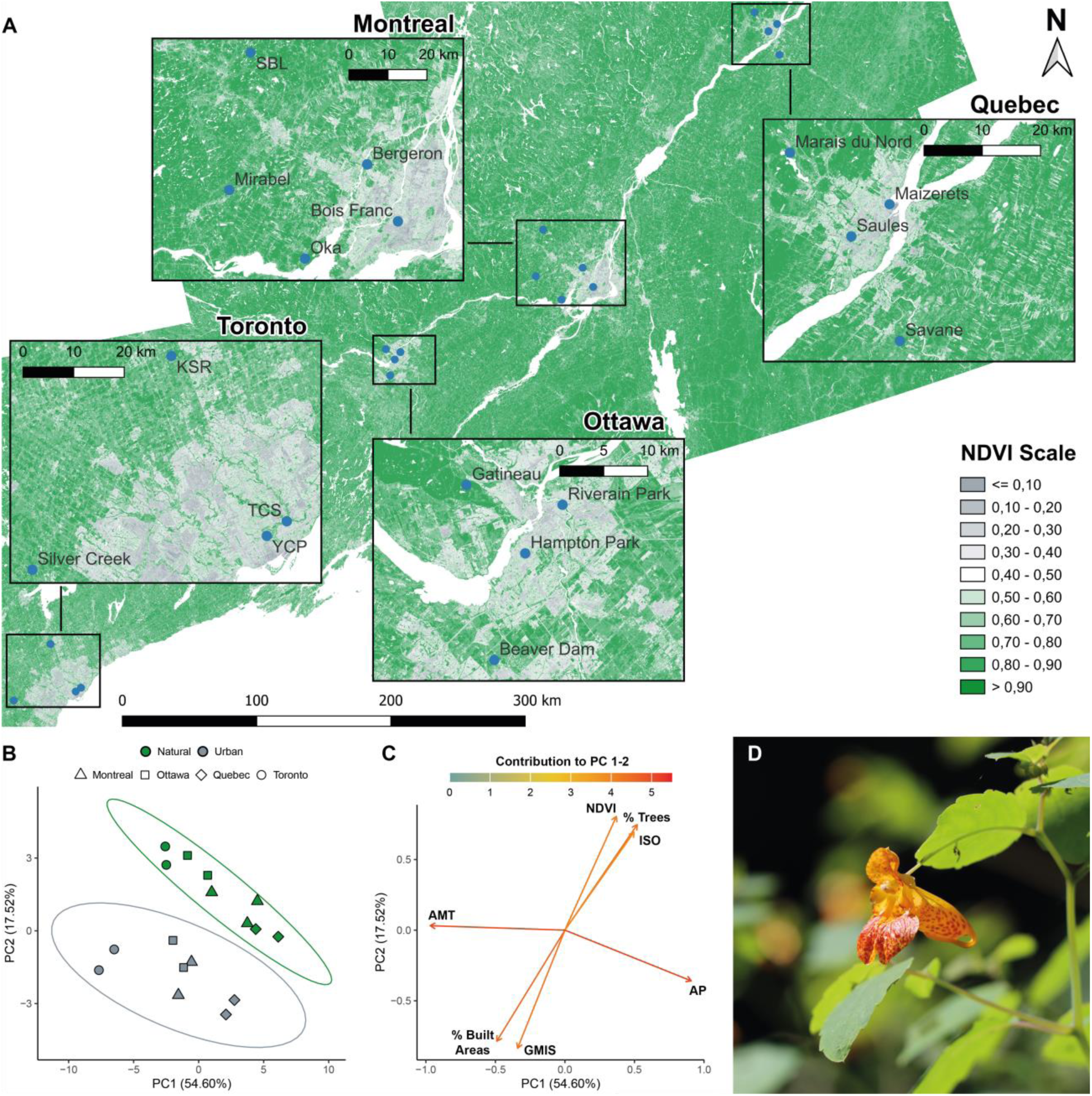
Study sites and system. Flowers from populations of *Impatiens capensis* were sampled in four regions in 2021 and 2022. (A) Normalized-Difference Vegetation Index (NDVI) map showing the sampled populations; high NDVI values (green) indicate more important vegetation cover. (B) Populations and (C) variables plots of a Principal Component Analysis (PCA) conducted on the mean values of environmental variables from a 500m radius circle around each population. Only the most contributing variables are represented: annual mean temperature (AMT), annual precipitation (AP), isothermality (ISO), NDVI, global man-made impervious surfaces (GMIS), percentage of trees (% Trees) and percentage of built areas (% Built Areas). Ellipses represent 95% confidence interval (CI). PCAs for 1000m and 2000m radius buffers are available in the Supplementary Material. (D) Flower of *Impatiens capensis* (credits: J. Burkiewicz).

In this study, we tested four hypotheses. First, the pollinators of *Impatiens capensis* differ in both their composition and sizes between urban and natural areas habitats. Second, because the pollinators of *I. capensis* must pass through the large tubular posterior sepal to access the nectar produced in the spur (Fig. 1), and because the reproductive organs are situated at the top entrance of the sepal, we expect sepal size to be correlated with the size of pollinators, and thus to differ between habitats, to better fit the local pollinator communities and ensure an effective pollen transfer. Third, we expect the corolla surface to be larger in urban habitats due to its role in pollinator attraction. This is based on previous studies that have shown that increased competition for floral resources (Hennig & Ghazoul, 2011) and reduced pollen dispersal in urban habitats (Rivkin et al., 2020) should result in stronger pollen limitation (Zink et al., 2024). Consequently, selection should favor larger floral displays in pollinator-limited populations to attract more pollinators (Geslin et al., 2013; Thomann et al., 2013). Fourth, sepal shape will differ between urban and natural habitats to maximize pollen transfer, but not corolla shape, as size appears more important than shape for pollinator attraction.

To test these hypotheses, we performed both field investigations and a greenhouse common garden experiment to control for environmental effects. We studied populations of *I. capensis* growing in sunlight to focus on the evolutionary responses of their animal-pollinated flowers. We considered the phenotypes measured in the common garden as the genetic components of the corresponding traits, whereas the phenotypes measured in the field corresponded to joint genetic and plastic components of the traits. We quantified flower shapes and sizes using geometric morphometrics, and we characterized pollinator communities for a total of 16 populations, paired by habitat – urban versus natural – in and around four large Canadian cities (“regions” hereafter): Toronto, Ottawa, Montreal, and Quebec (Fig. 1A).

## Materials and Methods

### Population selection and characterization

To test whether urbanization impacted the size and shape of *I. capensis* flowers, we compared urban and natural populations of four large Canadian metropolitan areas (+500,000 inhabitants). We selected two pairs of urban and natural populations with more than 100 individuals growing in open canopies in an area of 75 km around the city center of each region, for a total of 16 populations. Urban populations were selected to have high proportions of impervious surfaces, while natural populations were selected to have fewer impervious surfaces, more vegetation and few crop fields around them. One natural population selected in 2021 in the Toronto region produced too few chasmogamous flowers and was abandoned (only 15 populations were sampled in 2021). It was replaced with the Silver Creek population in 2022 (Table S1). The Oka natural population in the Montreal region in 2021 was replaced by the Mirabel population in 2022 because of access issues (Table S1).

To confirm that urban and natural sites were clearly distinct across regions, we characterized each population with the Normalized Difference Vegetation Index (NDVI) and the percentage of land use/land cover using satellite images from Sentinel-2 (resolution 10m), and the mean Global Man-made Impervious Surface (GMIS) (Brown de Colstoun et al., 2017) using Landsat 7 data (resolution 30m). We measured the mean NDVI, GMIS, and land use percentages for each population with buffer zones of 500 m, 1000 m, and 2000 m radius centered on them. We also characterized each population using bioclimatic variables. We interpolated values of temperature (minimum and maximum, mean, mean minimum and maximum, mean dew), precipitation (total, mean relative humidity, number of wet and dry days), solar radiation, and frost-related metrics (number of days with and without) using BIOSIM 11 (Régnière & Bolstad, 1994), and computed the monthly summary for each population between 2010 and 2022 to obtain average bioclimatic conditions across several years for each population. These values were used to compute worldclim bioclimatic variables using the QBMS package (Al-Shamaa, 2021) for all populations in R version 4.3.3 (R Core Team, 2024). To compare the populations, we ran a scaled principal component analysis (PCA) including all above abiotic variables.

### Field sampling

In both 2021 and 2022, we visited each population to photograph flowers, collect pollinators, and record flower visitors with cameras, and a second visit allowed us to collect pollinators and record flower visitors again. Both visits were done during peak flowering (between August 1^st^ and September 10^th^). In each population, we photographed 50 flowers in profile and front view from randomly selected plants with either a Canon T2i with a EF-S 60mm macro lens (1:2.8 USM, Canon Inc., Tokyo, Japan) or a Canon 90D with a EF 100mm macro lens (1:2.8 L IS USM, Canon Inc., Tokyo, Japan), placing a ruler beside flowers. We also recorded flower visitors of at least 5 different flowers for one hour using two GoPro Hero 9 cameras (GoPro, San Mateo, California, USA) placed 2 to 50 m apart. The camera was positioned 30 to 50 cm from the flowers, and recordings were done in 4K (3820 x 2160), with 30 frames per second. In 2021, we also sometimes recorded pollinator visits using a Canon T2i with a EF-S 60mm macro lens in 1080p (1920 x 1080) at 30 frames per second.

After the cameras stopped recording, two people collected insect pollinators of *I. capensis* for further identification and measurements during 20-minute sessions, using entomological nets while standing in front of, or walking along a stand of *I. capensis*. Pollinators were caught if they touched the reproductive parts of the flower and were then placed in a 50 mL tube in a cooler and frozen at the end of the day. All pollinator observations and captures were conducted in sunny conditions, with a minimum temperature of 15°C and low to no wind.

### Common garden experiment

During the summer of 2021, we collected one seed from at least 30 plants in each population for a common garden experiment. A few days after collection, seeds were stored in Petri dishes on moist filter paper, closed with parafilm, and maintained at 4°C until germination (ca. 4-5 months later). Seedlings were transferred to 2-inch square pots containing BM6 multipurpose soil (Berger, Saint-Modeste, Quebec, Canada) and watered every two days from the bottom of the tray, alternating between clear water and fertilized water (15-5-15 NPK fertilizer from Monday to Thursday and 19-2-19 NPK from Friday to Sunday). When the plants reached 15 cm in height, they were transferred to 8.5 inch round pots with BM6 soil. They were watered and fertilized every two days, and every day from March 1^st^. Plants were grown with 16 hours of light, with day and night temperatures of 22°C and 18°C, respectively. When plants started flowering, we photographed the front and profile views of one random flower on each plant for phenotyping, using a Canon T2i with an EF-S 60mm macro lens, including a ruler on each picture (Table S1). The common garden experiment aimed to remove the environmental effects on the plants’ phenotypes, but because it consisted of the progeny of the natural plants, maternal effects might be expressed.

### Pollinator identification and measurement

Frozen pollinators were pinned on foam blocks using needles and dried at room temperature for a week. Apart from one Calliphoridae, one Muscidae, and one wasp specimen (the last being identified at the genus level) pollinators were identified at the species level. Pollinators were deposited at the Ouellet-Robert entomological collection of the Université de Montréal (Burkiewicz, 2025). We used the inter-tegular distance (ITD) as a proxy of pollinator size, as it is a good indicator of insects’ dry mass and flight capability (Goulson, 2010; Greenleaf et al., 2007). We measured it by photographing each pollinator under a Zeiss microscope equipped with an AxioCam HRc (Discovery V20) camera and using the software ZENblue v2.5 Pro (Carl Zeiss, Oberkochen, Germany).

### Pollinator observations

On the camera recordings, we identified between 5 and 10 open flowers from which we could easily distinguish a pollination event. An animal was considered a pollinator when it touched the reproductive parts of the flowers. Videos were watched at 2 × speed. When an animal was observed, videos were slowed down (0.12 – 0.5 ×) to score pollination events. For each pollinator, we recorded the time of the bout, the species or genus and family, the number of flowers observed in this particular recording, and the number of flowers visited. We identified bumblebees at the genus level, other bees at the species level, and wasps and flies at the family level. When we could not determine the species or genus, we identified taxa at the family level.

We calculated the total observation effort as the sum over all observation units (eg., one camera recording) of the number of hours observed times the number of flowers observed per unit. The pollinator visitation rate was the number of flowers pollinated over the total observation effort. Finally, the relative frequency of each pollinator taxon was the number of pollination events observed for that taxon divided by the total number of pollination events, either at the population level or over all populations.

### Flower shape and size characterization

We used geometric morphometrics to characterize the flower shape from flower pictures taken in profile and front views, and measure the posterior sepal size and corolla surface (Collyer et al., 2015; Klingenberg, 2010; Mitteroecker & Gunz, 2009). We used tpsDig v2.32 (Rohlf, 2021) to digitize specimens. We used a similar approach to that of Faure et al. (2023) to extract flower shape (Fig 5A, D), and details on landmarks and semilandmarks positioning are available in Supplementary Materials. Flower size was estimated from the landmarks and semi-landmarks that delimited the contour of the sepal and the corolla (Fig 3A, E); more details can be found in the Supplementary Materials. To compare flower shape between populations we performed a General Procrustes Analysis (GPA) with the *gpagen* function of the geomorph package (Adams, 2020), minimizing Procrustes distance between individuals (Klingenberg, 2010). As in Faure et al. (2023), semilandmarks were treated as landmarks in the GPA because the high variability of the spur and petals’ curvature caused misalignments when treated as semilandmarks. We manually checked for outlier shapes to ensure these were not due to improper landmark positioning.

To assess the technical error in landmark positioning, we placed landmarks on duplicate flowers in some populations (four in front view, five in profile view). The technical error estimated using a Procrustes ANOVA with 999 permutations (*procDlm* function of the geomorph package (Adams, 2020)) was 0.02% for the profile pictures and 2.08% for the front pictures.

### Statistical analyses

All analyses were performed in R version 4.3.3 (R Core Team, 2024) using the car (Fox et al., 2001), emmeans (Lenth, 2017), lme4 (Bates et al., 2003), geomorph (Adams, 2020), and vegan (Oksanen et al., 2025) packages. We visually inspected the normality and homoscedasticity of residuals, indicating if these assumptions were unmet and how we addressed these cases.

#### Pollinator abundance

We evaluated whether pollinator communities differed between urban and natural habitats. Because observation data and collected specimens gave consistent results (Fig. 2A, Table S2), we used only collected specimens for statistical analysis, as most insects were identified at the species level. Redundancy analysis (RDA) was used to analyze community data rather than canonical correspondence analysis (CCA) because in the latter, rare species contribute importantly to the chi-square distance (Legendre and Legendre 2012), and we were more interested in differences in the main pollinators. We used a Hellinger transformation of the data to account for shared absences among sites (Legendre and Legendre 2012). We tested whether regions (cities) or habitats (urban/natural) could explain variation in pollinator abundance. Partial RDA was used to test each factor while removing the effect of the other.

**Fig. 2.**
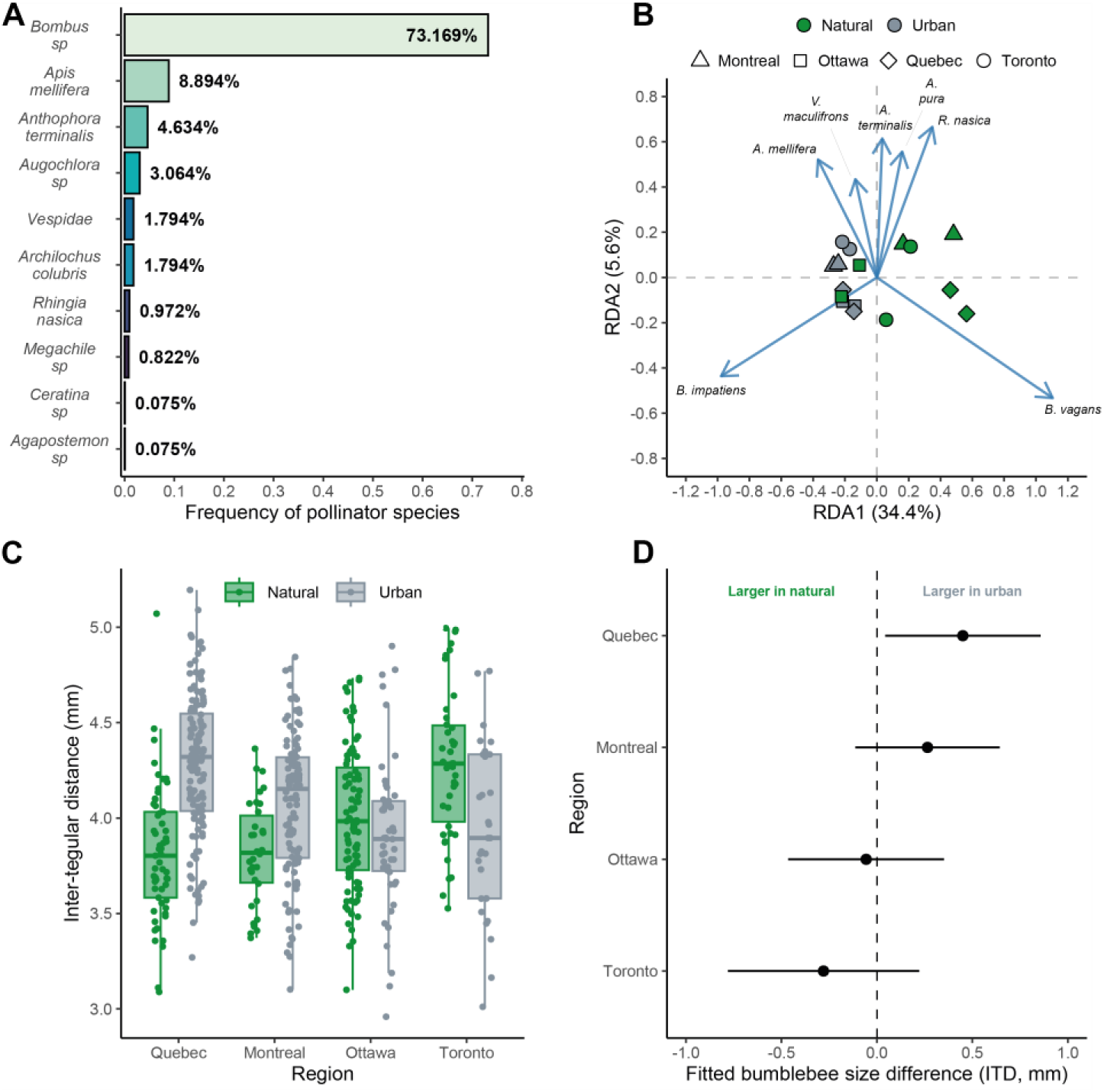
Characterization of the *I. capensis* pollinator community and size of bumblebees. (A) Relative frequencies of pollinator species computed from video observation data, although frequencies are similar to those obtained with collected specimens (Table S2). (B) Redundancy analysis of pollinator abundance (collected specimens) by habitat and region for the 2022 field season. Only the most abundant species were represented on the graph for clarity purposes. The redundancy analysis on combined field seasons (2021 and 2022) is available in Fig. S2. (C) Boxplots of bumblebee inter-tegular distances by habitat and region, averaged over the two field seasons. Dots represent individual bumblebee measurements; sample sizes are available in Table S4. (D) Estimated marginal means contrasts of bumblebee size between urban and natural habitats for each region from the linear mixed model and associated 95% confidence intervals. Measures of bumblebee size include all *Bombus sp.* collected.

#### Pollinator visitation rate

We assessed whether urbanization influenced pollinator visitation rates. We used region, habitat and their interaction as fixed factors in an ANOVA to explain the variation in pollinator visitation rate, assuming a Gaussian distribution. Significance was tested using the *Anova* function from the car package and type II sum of squares.

#### Bumblebee size and flower size

We tested whether urbanization affected the sizes (ITD) of bumblebees (all species combined), the main pollinator of *I. capensis*, and of flowers (sepal size and corolla surface). We built a linear mixed model that included Year (when applicable), Region, Habitat, and their interactions as fixed effects, including a population random effect and allowing slopes to vary with region, habitat, and their interaction:

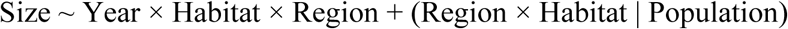

We first selected the best variance structure by fitting and comparing models with random effects of descending order of complexity and all fixed effects. Models were fitted with Restricted Maximum Likelihood (REML) using the *lmer* function from the lme4 R package and the best variance structure was the one with the lowest AIC (Zuur et al., 2009). Singular models and models that did not converge were excluded from the comparisons. The best variance structure for the three response variables was a simple population random effect, independent of region or habitat:

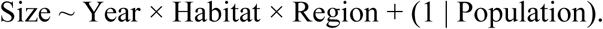

Using this variance structure, we then tested fixed effects and their interactions using type II Wald tests (*Anova* function from the car package). We computed estimated marginal means contrasts for urban/natural comparisons within regions within years using the Holm correction for multiple testing.

For flower size, we also tested if the sexual stage of the flowers (male or female) was correlated with the sepal size or the corolla surface, thus eliminating a potential confounding effect. The sexual stage was weakly associated (R^2^_adj_ <1%) with sepal size and corolla surface in the field. To further test the robustness of the results to the sexual stage, we compared the results of models that included or excluded female flowers (12.2% of the data) and found consistent results for both analyses (Table S6), indicating that our results were not affected by the sexual stage of the flower.

#### Plasticity of flower size

We tested whether flower size was affected by phenotypic plasticity and if this effect was influenced by urbanization by comparing the flower size measured in the field to that measured in the common garden. We first analyzed the data from the 2021 field season versus the common garden experiment to compare the phenotype of the parental population and the phenotype of the offspring generation (the parentage of which is unknown), showing both genetic and plastic responses to environmental change. We then analyzed the data from the 2022 field season versus the common garden experiment to compare the phenotypes of kin, representing the plastic response to environmental change.

For each comparison, we built a linear mixed model to analyze variation in flower size with Environment (either field or common garden), Habitat (either urban or natural), and their interaction as fixed effects, including a population random effect, allowing slopes to vary with environment, habitat, and their interaction. We used the same procedure mentioned above to select the best variance structure. The best model included a population random effect that changed the intercept and the slopes were allowed to vary with Environment:

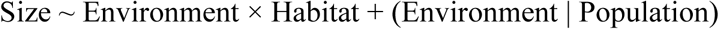

We then tested fixed effects and interactions using type II Wald tests using the *Anova* function from the car package.

#### Correlation between bumblebee size and sepal size, and between pollinator visitation rates and corolla surface

We tested whether variation in mean population sepal sizes (or corolla surfaces) was correlated to the variation in mean population bumblebee sizes (ITD) (or mean population visitation rates). We used the sepal sizes and corolla surfaces measured in the common garden because they correspond to the genetic component of the trait, and analyzed their variation with an analysis of covariance with habitat and ITD (or the average pollinator visitation rates) and their interaction as fixed factors:

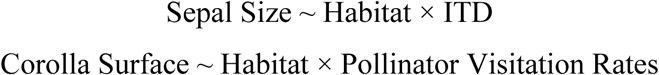

Significance was tested using the *Anova* function of the car package and type II sum of squares as above.

#### Flower shape

We used a Procrustes ANOVA (equivalent to a PERMANOVA) with the *procD.lm* function from the geomorph package to test whether flower shape was influenced by size (allometry effect), year (when applicable), sexual stage of the flower (male vs female), region, and habitat, including all possible interactions between these variables:

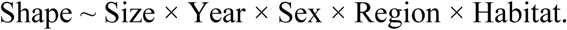

Because Procrustes ANOVA cannot include random factors, we followed (Collyer et al., 2015) and used the interaction of habitat and population as the error term to test for fixed effects on interpopulation variation. We removed all non-significant interactions, except the one between habitat and region, which we were interested in, and ran a Procrustes ANOVA with the remaining terms. We used type 1 Procrustes ANOVA, ordering factors in a way that we tested for the effect of habitat and the interaction between habitat and region after all other factors.

## Results

### Urban populations were environmentally distinct from natural populations

Urban and natural populations, across the four regions studied, were characterized by different abiotic environments, showing our ability to discriminate those environments in the field and justifying the use of the “habitat” categorical variable in all analyses that quantify the effects of urbanization (Fig. 1 and Fig. S1). The natural populations were associated with higher tree cover and greater vegetation index (NDVI), whereas urban populations were associated with higher proportions of impervious surfaces and built areas, as well as lower isothermality (day-to-night temperature variation compared to annual oscillations) due to a heat island effect (Fig. 1B, C and Fig. S1).

### Urbanization modifies the pollinator communities of *I. capensis*

Bumblebees (*Bombus* spp.) were the most frequent pollinators of *I. capensis*, representing 73% of all pollination events recorded on video (Fig. 2A). *Bombus impatiens* was the most collected species (62.4%, Table S2), followed by *Bombus vagans* (12.7%, Table S2). *Apis mellifera* represented 6.4% of the specimens sampled (Table S2) but primarily collected pollen and may not be a reliable pollinator, as stigmas are not receptive when anthers are mature. *Anthophora terminalis*, *Vespidae*, and *Augochlora pura* together represented 12.5% of the specimens collected (Table S2), and these less common and less cosmopolitan pollinators did not make contact as consistently with the reproductive structures as bumblebees, according to video recordings.

Pollinator communities of *I. capensis* differed between urban and natural habitats (R^2^_adj_ = 0.264, P = 0.013; Fig. 2B, Fig. S2). Urban habitats were more associated with *Bombus impatiens, Vespula maculifrons,* and *Apis mellifera,* while natural habitats were more associated with *Bombus vagans*, *Augochlora pura* and *Rhingia nasica* (Fig. 2B, Fig. S2). Among these pollinators, bumblebees were the most consistently found across all populations, and as such, they best characterized urban and natural pollinator communities of *I. capensis.* We also found that the effect of urbanization on pollinator visitation rates varied among regions (Region × Habitat: F_(3, 23)_ = 3.675, P = 0.026; Table S3A, B).

Finally, we found that the effect of urbanization on bumblebee size varied among regions (Region × Habitat: **χ**^2^_(3)_ = 8.826, P = 0.031; Fig. 2C, D, Table S5A), with Quebec showing the strongest effect (11.6% increase from natural to urban populations, Fig. 2C, D, Table S5B).

### Urbanization affects the genetic component of sepal size

Sepal size was the same for urban and natural populations in the field (**χ**^2^_(1)_ = 0.239, P = 0.624; Fig. S3, Table S6C). However, in the common garden experiment, sepals from urban populations were on average 7.6% larger than those in natural populations (**χ**^2^_(1)_ = 8.564, P = 0.003; Table S7A), though this effect varied among regions (Region × Habitat: **χ**^2^_(3)_ = 8.329, P = 0.039; Table S7A), with Quebec populations showing the largest difference (20.3% increase, t_(7.29)_ = 3.700, P = 0.028; Fig. 3B, C, Table S7B). Together, these results suggest that phenotypic plasticity in the field could hide the genetic differences in sepal size between habitats. To further investigate this phenotypic plasticity, we analyzed the mean response of sepal size per habitat to changes in the environment (field vs. common garden). Compared to the common garden experiment, the sepal size measured in the field was smaller in urban populations and larger in natural populations, thus completely masking the existing genetic differences in sepal size between habitats (Environment × Habitat, 2021: **χ**^2^_(1)_ = 4.796, P = 0.028; 2022: **χ**^2^_(1)_ = 3.706, P = 0.054; Fig. 3D, Fig. S4A, Table S8).

**Fig. 3.**
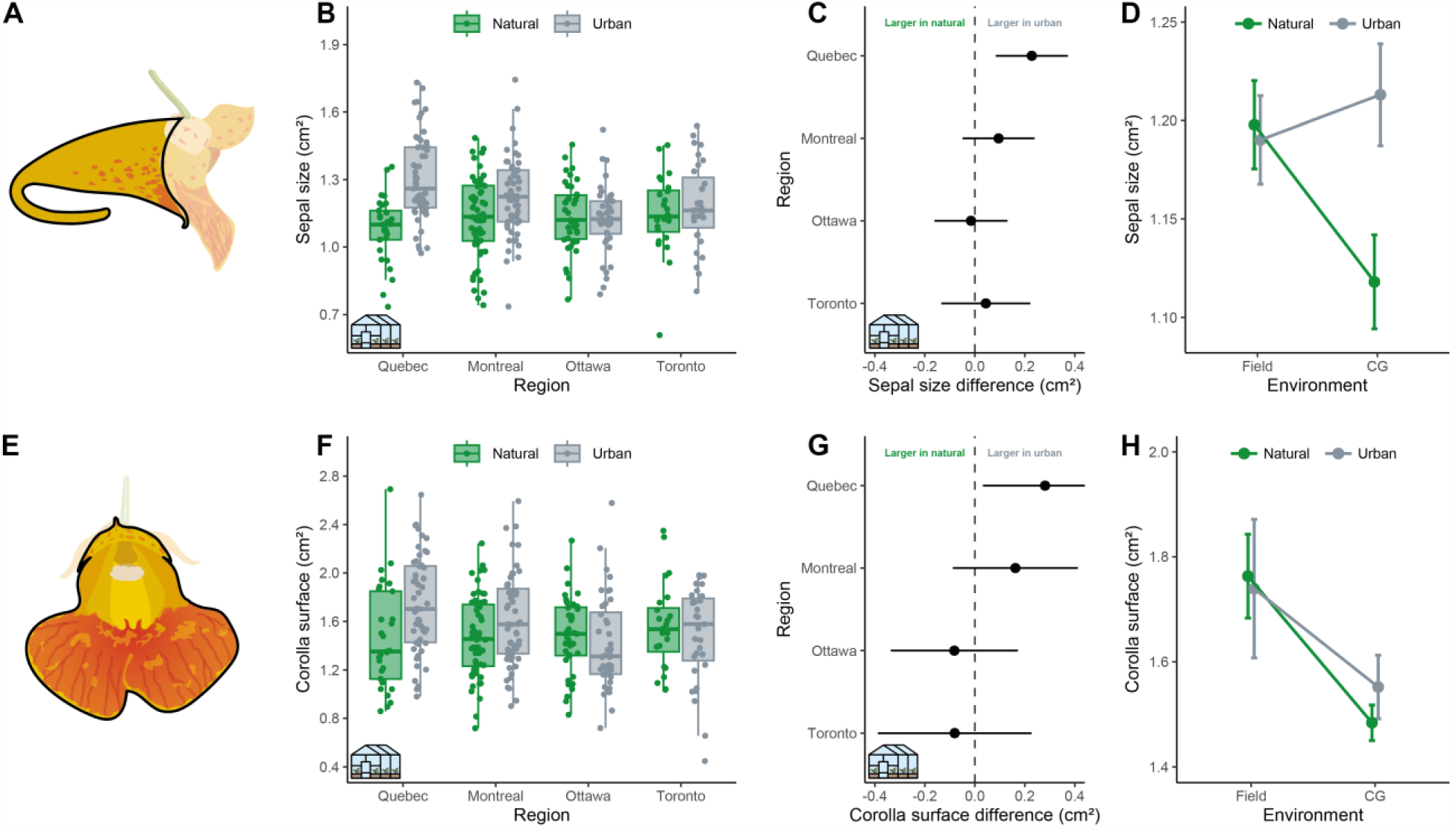
Sepal size (A-D) and corolla surface (E-H) comparisons between habitats. (A) Drawing showing the measure used to estimate the sepal size of *Impatiens capensis* in black outline. (B) Boxplots of the sepal size measured in the common garden (as indicated by the greenhouse icon) with dots representing individual flowers. (C) Estimated marginal means contrasts of the sepal size between urban and natural habitats per region and associated 95% confidence intervals in the common garden experiment. (D) Mean response of sepal size, grouped by habitats, to environmental change for the 2021 field season. Error bars represent standard errors of the mean. Results with two field seasons combined are available in Fig. S5A. (E) Drawing showing the measure used to estimate the corolla surface of *Impatiens capensis* in black outline. (F) Boxplots of the corolla surface measured in the common garden with dots representing individuals. (G) Estimated marginal means contrasts of the corolla surface between urban and natural habitats per region and associated 95% confidence intervals in the common garden experiment. (H) Mean response of corolla surface, grouped by habitats, to environmental change for the 2021 field season. Error bars represent standard errors of the mean. Results with two field seasons combined are available in Fig. S5B. Credits for the drawings: J. Lafond.

We found that the genetic component of the mean sepal size per population was positively correlated with the mean population bumblebee size (F_(1,11)_ = 6.531, P = 0.026, R^2^_adj_ = 0.527; Fig. 4, Table S9).

**Fig. 4.**
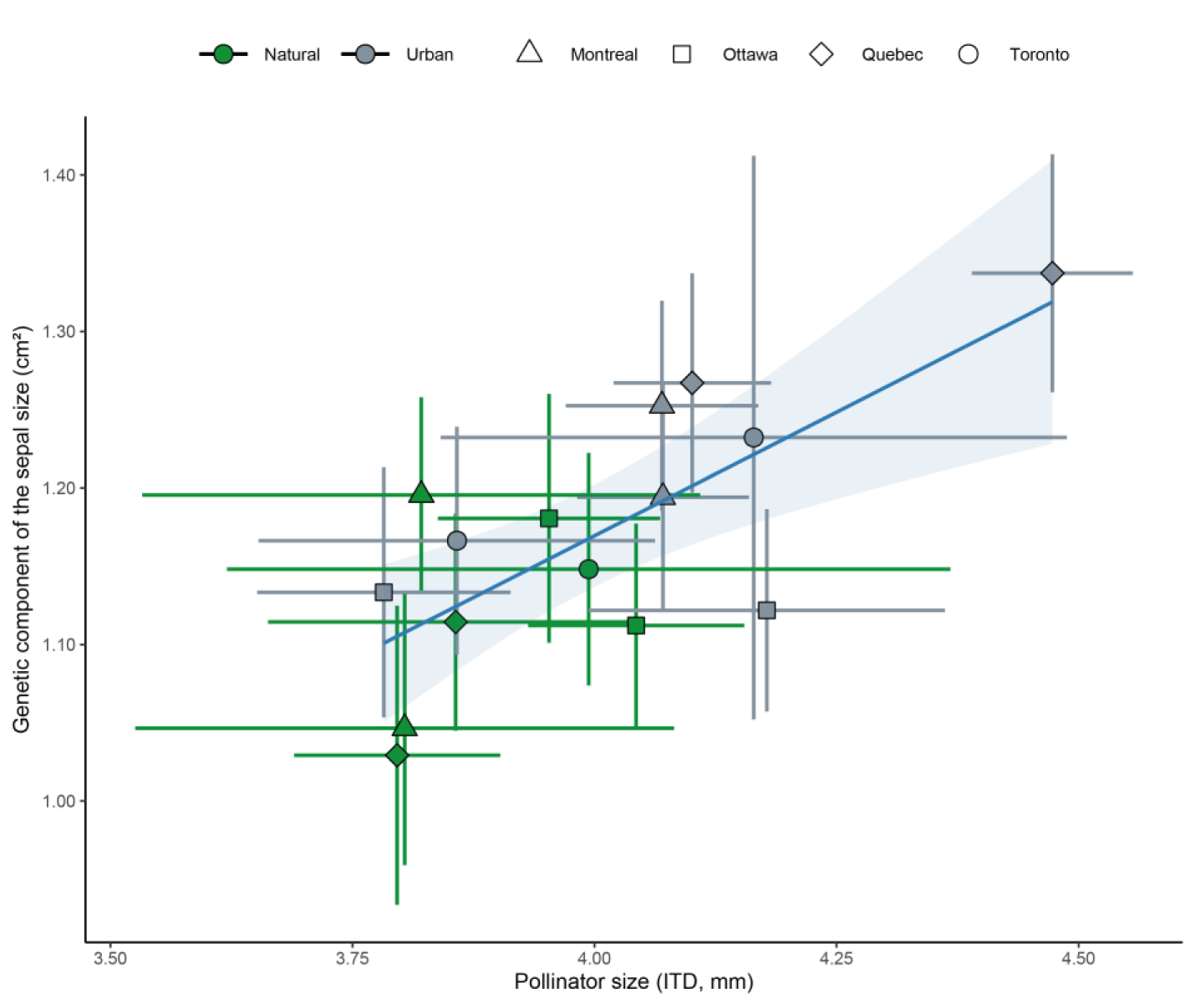
Correlation between the genetic component of the sepal size and bumblebee inter-tegular distances. The points represent the population mean of the sepal size in the common garden experiment and the population mean bumblebee size averaged across the 2021 and 2022 field seasons. Error bars for the points indicate the 95% confidence intervals of the means. The line shows the significant positive linear relationship with its standard error. Measures of bumblebee size include all *Bombus sp.* collected.

### Urbanization has a limited effect on corolla surface, which is strongly affected by phenotypic plasticity

In the field, corolla surface was affected by habitat, but only in certain regions and certain years, suggesting phenotypic plasticity (Table S6C). Under controlled conditions, the effect of urbanization on corolla surface varied among regions (Region × Habitat: **χ**^2^_(3)_ = 8.136, P = 0.043; Fig. 3F, G, Table S7A). The corolla surface was 19% larger in the field compared to the common garden, regardless of habitat, confirming strong phenotypic plasticity for this trait (2021: **χ**^2^_(1)_ = 8.859, P = 0.002; 2022: **χ**^2^_(1)_ = 36.761, P < 0.001; Fig. 3H, Fig. S4B, Table S8). The genetic component of the corolla surface was positively correlated with pollinator visitation rates in urban populations, but not in natural populations (Pollinator Visitation Rate × Habitat: F_(1,11)_ = 14.851, P = 0.002; Fig. S7, Table S10)

### Urbanization affects the sepal shape of *I. capensis* through plastic responses

The spur of urban flowers was shorter and less curved compared to the ones in natural habitats in the field (F_(1,1518)_ = 3.751, P = 0.019; Fig. 5B, C, Table S11A), but not in controlled conditions (F_(1,315)_ = 0.858, P = 0.502; Fig. 5B, C, Table S11A). In contrast, corolla shape was similar in urban and natural habitats, both in the common garden (F_(1,314)_ = 0.436, P = 0.837; Fig. 5E, Table S11A) and in the field (F_(1,1509)_ = 0.886, P = 0.472; Fig. 5F, Table S11A). Corolla shape of flowers from the field populations also differed from those raised in the common garden (Environment: F_(1,1839)_ = 6.198, P = 0.002; Fig. 5E, F, Table S11B), further supporting the effect of phenotypic plasticity on flowers. The shape of the corolla in controlled conditions suggests that they are less open than those in the field (Fig. 5E, F).

**Fig. 5.**
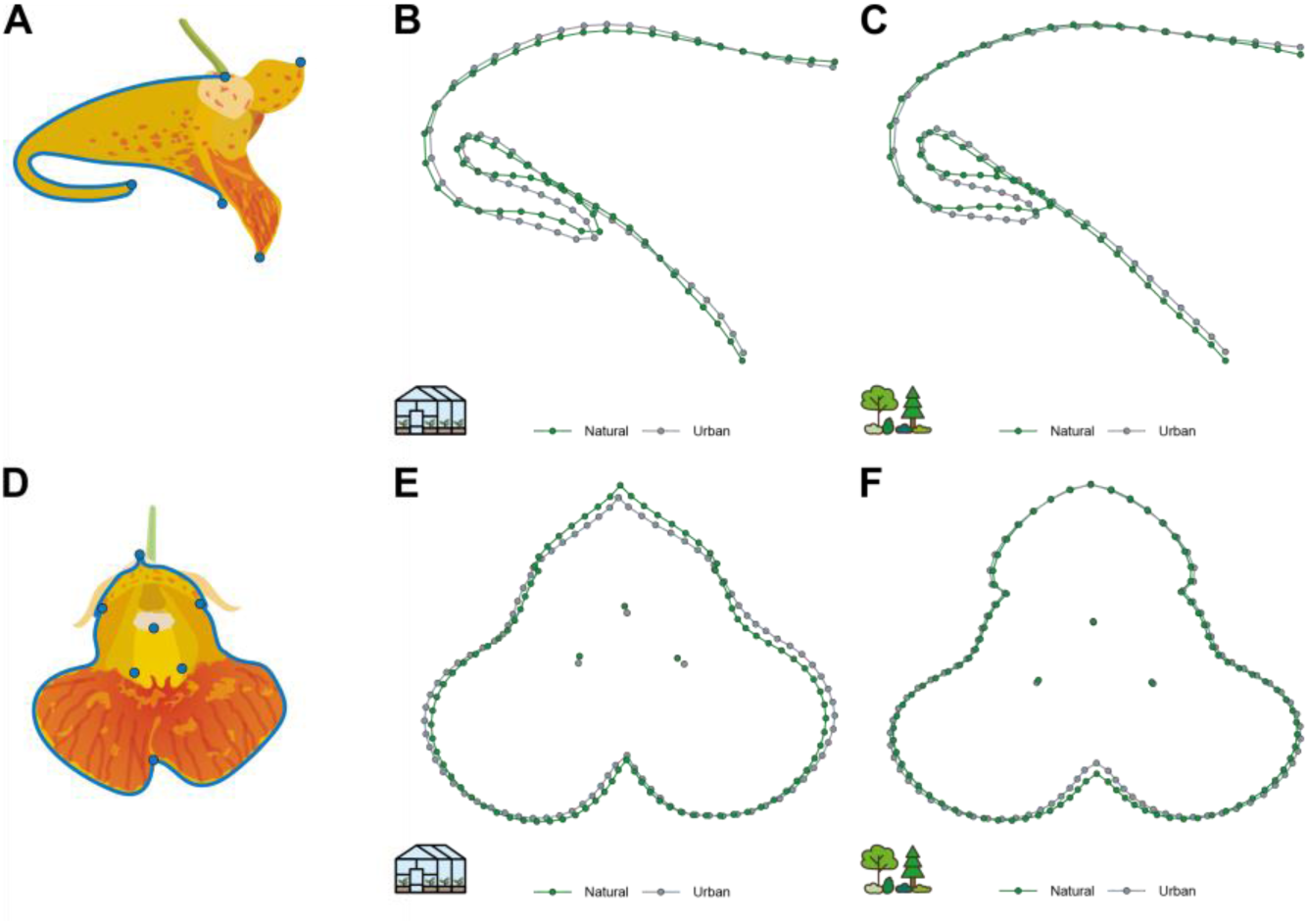
Predicted posterior sepal and corolla shape comparison between habitats. (A) Drawing showing the landmarks (blue dots) and semilandmarks (blue curves) position on the posterior sepal of *Impatiens capensis*. (B) Predicted shape of the posterior sepal in urban and natural habitats for the common garden experiment. (C) Predicted shape of the posterior sepal in urban and natural habitats for the 2021 field season. (D) Drawing showing the landmarks (blue dots) and semilandmarks (blue curves) position on the corolla of *Impatiens capensis*. (E) Predicted corolla shape in urban and natural habitats for the common garden experiment. (F) Predicted corolla shape in urban and natural habitats for the 2021 field season.

## Discussion

The combination of fieldwork and common garden experiments across four cities in southeastern Canada revealed that urbanization altered the pollinator communities of *I. capensis* and its flower morphology. Different species of pollinators were associated with urban and natural populations of *Impatiens capensis.* Moreover, urbanization affected bumblebee size, the main pollinator, differently among regions, a result explained by both intraspecific and interspecific differences. These differences in pollinator community and size support our initial hypothesis. The genetic component of sepal size was larger overall in urban areas, despite variation among cities. In addition, and supporting our second hypothesis, we found that the genetic component of sepal size was positively correlated with bumblebee size across populations, in both urban and natural habitats. This suggests that, despite substantial phenotypic plasticity in the field that sometimes masked the genetic determination of sepal size, as well as the existence of discrepancies among regions, we may have found an explanation for the sepal size variation of *I. capensis* across urbanization gradients, where sepal size has to fit the size of its bumblebee pollinators to maximize pollen transfer.

Urbanization affected the genetic component of the corolla surface differently among regions, with corollas being significantly larger in Quebec’s urban areas, partly supporting our third hypothesis. However, corolla surface was positively correlated with pollinator visitation rates in urban populations (Fig. S7), contrary to our expectation that larger corollas should be favored in habitats with reduced pollination services. This finding does not preclude the expected negative relationship between floral display and pollen limitation as our analysis was limited to a single component of floral display, corolla surface, excluding the effect of flower number, and did not account for pollen transfer effectiveness and pollen quality in the estimation of pollen limitation. Trade-offs may also play a role; for example, limited resources could constrain investment in both flower size and number, such that increased flower number compensates for smaller corollas in overall display. Pollinator visitation rates to *I. capensis* may also be driven by the entire plant community through both competitive and facilitated interactions with co-flowering species. For instance, the presence of the large-flowered invasive *Impatiens glandulifera* has been associated with selection for reduced corolla height in *I. capensis* (Beans & Roach, 2015).

For both sepal size and corolla surface, the impact of urbanization varied among cities, but it aligned with the impact of urbanization on pollinator communities. For example, pollinators were the largest in urban populations of Quebec, where the genetic component of flower size was the most impacted by urbanization. Previous studies on flower evolution in urban areas have focused on phenotypic changes associated with pollinator availability and pollen movement (Irwin et al., 2018; Ushimaru et al., 2014), but they left aside a more thorough characterization of pollinator communities. Here, we show that characterizing the pollinator community helps to understand the floral evolution in an urbanization context.

Urbanization had a limited effect on flower shape, partly supporting our fourth hypothesis. In the field, urban sepals had shorter and less curved nectar spurs, which partially supports the results of Faure et al. (2023), who found that urban nectar spurs were less curved in the region of Montreal. However, the lack of difference in the common garden experiment suggests that urbanization affected sepal shape only through phenotypic plasticity, although the statistical power to detect a genetic response was small for the common garden experiment (Table S1). In contrast, corolla shape was not affected by urbanization in both field and common garden conditions. Overall, flower shape—excluding size—shows no evidence of genetic adaptation and seems to be strongly influenced by environmental factors. These findings may seem surprising, as spur curvature, a component we characterize through sepal shape, has been linked to improved pollen removal by hummingbirds (Travers et al., 2003) and longer visit durations by bees (Young, 2008). However, in our populations, bumblebees were the main pollinators, and hummingbirds were rare, potentially limiting their role as selective agents. Finally, there is no reason to believe that visit duration by bumblebees differs between urban and natural habitats, so the lack of difference in our common garden experiment appears logical.

Phenotypic plasticity can be adaptive or maladaptive depending on whether it moves phenotypes towards or away from the fitness optimum of populations. It can also decrease the efficiency of natural selection as the phenotypes with the highest fitness represent both the genetic and plastic components of the corresponding traits. We found that for the sepal size of *I. capensis*, phenotypic plasticity masked the genetic differences detected in the common garden experiment (Fig. 3D; Fig S4), suggesting a counter-gradient variation in which plasticity typically modifies the phenotype in a direction opposite to natural selection (Conover & Schultz, 1995). Assuming a similar plastic response for all genotypes, such as larger flowers in the common garden are also larger in the field conditions, we infer that the sepal size genetically respond to its pollinator community. Therefore, bumblebee selection for sepal size along an urbanization gradient appears to be a case of compensatory adaptive change in which selection counteracts the homogenizing effect of phenotypic plasticity (Conover et al., 2009; Conover & Schultz, 1995; Hendry, 2016). Indeed, natural selection seems to favor larger flowers in some cities because of the presence of larger bumblebees, but plasticity masked the genetic differences in the field (Fig. 3D, Fig. S4). We also highlighted that the corolla size and shape were significantly affected by the environment through phenotypic plasticity, which is particularly evident from the observed differences between the common garden and field conditions. This plasticity could be caused by developmental constraints, weather conditions, and pollinator visits (by pushing the petals apart), and may at least partly explain the variation among years (Table S6C), and between field and controlled conditions. Our results demonstrate that it is critical to disentangle plastic from genetic responses to fully understand the causes of phenotypic changes and adequately interpret the evolutionary responses of organisms to environmental changes (de Villemereuil et al., 2016; Diamond & Martin, 2021; Lambert et al., 2021; Thompson et al., 2025).

Strong environmental changes associated with urbanization impact living organisms, their interactions, and their evolution. Although our knowledge of eco-evolutionary dynamics in these environments has increased in the last decade, integrative approaches that assess plastic and genetic components of traits while considering species interactions are still scarce, but they are necessary to understand rapid evolution to changing environments. Only with this will we be able to understand adaptation to urbanization and pave the way for conservation policies in these habitats.

## Acknowledgments

In Ottawa, sampling was allowed by the National Capital Commission with permits 23562 in 2021 and 24501 in 2022. In the Silver Creek Conservation Area, sampling was allowed with permit CL22/021. We thank the Koffler Scientific Reserve and staff for access, housing, and help during our sampling. We thank Real Collins and Shirley Dumais for granting access to their property in Mirabel. We thank Agiro for granting access to Marais du Nord for sampling, and Domaine de Maizerets for access to sample in the Maizerets population. We thank C. Fauteux, M. Vallée, O-L. Kuhn, A. Corkal, and M. Duguay for assisting with fieldwork and data analysis. We thank Guillaume Larocque for his help with the NDVI map. We thank Daniel Schoen for his comments on a previous version of this manuscript.

## Author contributions

Conceptualization: SJ, JB

Methodology: JB, SJ, CD

Investigation: JB, JC, SC, JL, EN, SJ

Visualization: JB, SJ, CD, JL

Funding acquisition: SJ

Project administration: JB, SJ

Writing – original draft: JB

Writing – review & editing: JB, JC, SC, JL, EN, CD, SJ

## Data and code availability

All code and data are available in figshare (private upon acceptance) and Canadensys (Burkiewicz, 2025)

## Supplementary Materials

### Supplementary Methods

#### Geometric morphometric

We identified five landmarks (homologous points) for the posterior sepal: 1) the tip of the upper petal, 2) the insertion of the pedicel on the perianth, 3) the extremity of the spur, 4) the lower extremity of the opening of the posterior sepal, and 5) the lower point of the lower petal closer to the camera (Fig. 5A). We also defined two curves with 30 semilandmarks each to capture the shape of the posterior sepal. The first curve followed the outline of the posterior sepal between landmarks 2 and 3 (dorsal curve, Fig. 5A), and the second one followed the outline of the posterior sepal between landmarks 3 and 4 (ventral curve, Fig. 5A).

We placed seven landmarks for the front views of the flowers: 1) the tip of the upper petal, 2) the intersection between the upper petal and the right lateral petal (left for landmark 6, which is symmetrical), 3) the right widest point of the corolla opening (left for landmark 5, which is symmetrical), 4) the intersection of the two lower petals or, if the two petals do not intersect, the center of a segment drawn between the uppermost part of each lower petal, and, 7) the lower point of the stamens or the stigma (Fig. 5D). Note that points 4 and 7 are better defined as semi-homologous landmarks. We also defined four semilandmarks curves to capture the shape of the corolla. The first and fourth followed the outline of the upper petal between landmarks 1 and 2, or 6 and 1, and each contained 10 points. The second and third contained 35 points and followed the lateral and lower petal outline between landmarks 2 and 4, or 4 and 6 (Fig. 5D). Curves were numbered clockwise starting from the top of the flower.

We removed flower pictures that were too blurry or for which critical features of the flowers were masked, preventing us from precisely locating landmarks or semilandmarks. For all curves, we removed the first and last semilandmarks because they were redundant with the landmarks. For profile pictures, we also removed landmarks 1 and 5 because they were redundant with the landmarks and semilandmarks of the front pictures and were sometimes challenging to locate precisely. Different persons placed the landmarks, but one person (JB) validated all points and curves of all flowers.

For the profile view, the sepal size corresponded to the area of a polygon created by concatenating all curves and landmarks 2, 3, and 4 (Fig. 3A, Fig. 5A). For the front view, the corolla surface corresponded to the area of a polygon created by concatenating all curves and landmarks 1, 2, 4 and 6 (Fig. 3E, Fig. 5D).

## Supplementary Figures

**Fig. S1.**
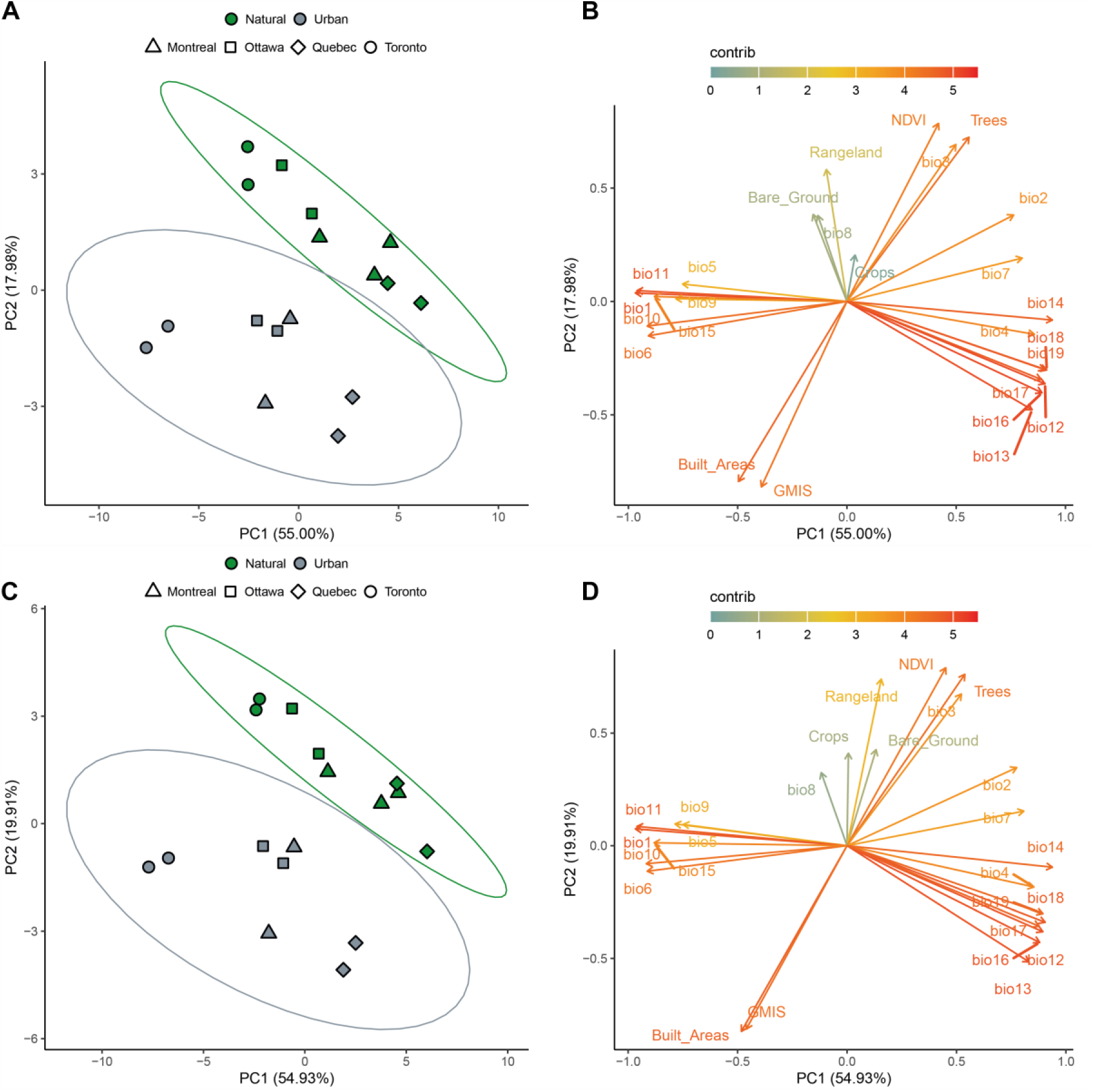
Population characteristics at different spatial scales. (A-B) Individual and variables plot of a Principal Component Analysis (PCA) on urbanization and bioclimatic variables computed using a circle of 1000m radius around each population. (C-D) Individual and variables plot of a Principal Component Analysis (PCA) on urbanization and bioclimatic variables computed using a circle of 2000m radius around each population. Urban and natural populations mainly differ in the percentage of impervious surfaces, trees and isothermality (BIO3) in both analyses, whereas cities vary according to temperatures and precipitations.

**Fig. S2.**
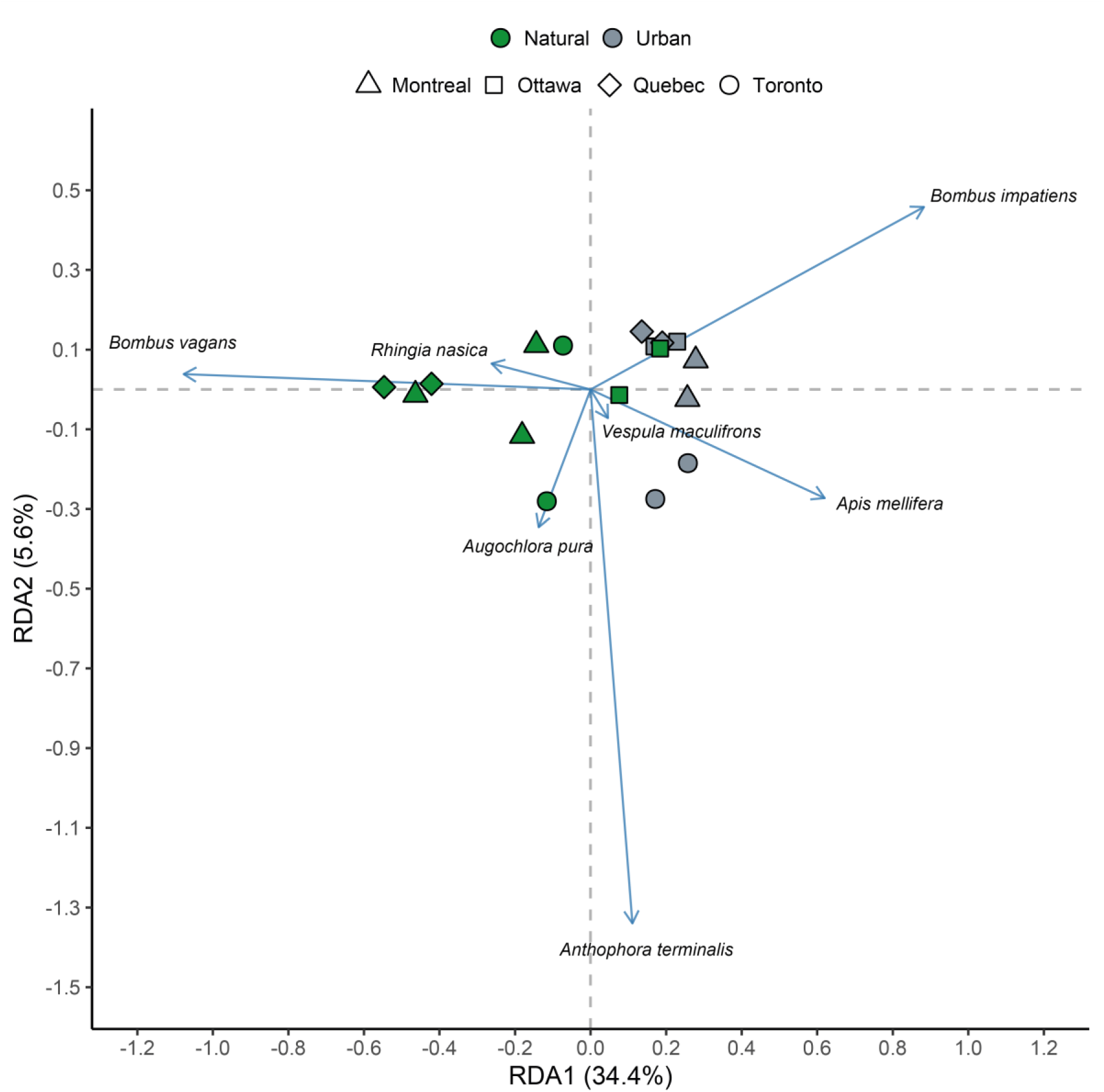
Redundancy analysis (RDA) evaluating the impact of urbanization on the *Impatiens capensis* pollinator community in 2021 and 2022. When both years are considered together, the main difference between urban and natural habitats is the bumblebee species pollinating *I. capensis*. *Anthophora terminalis* separates the Toronto region because it was overrepresented in this region in 2021.

**Fig. S3.**
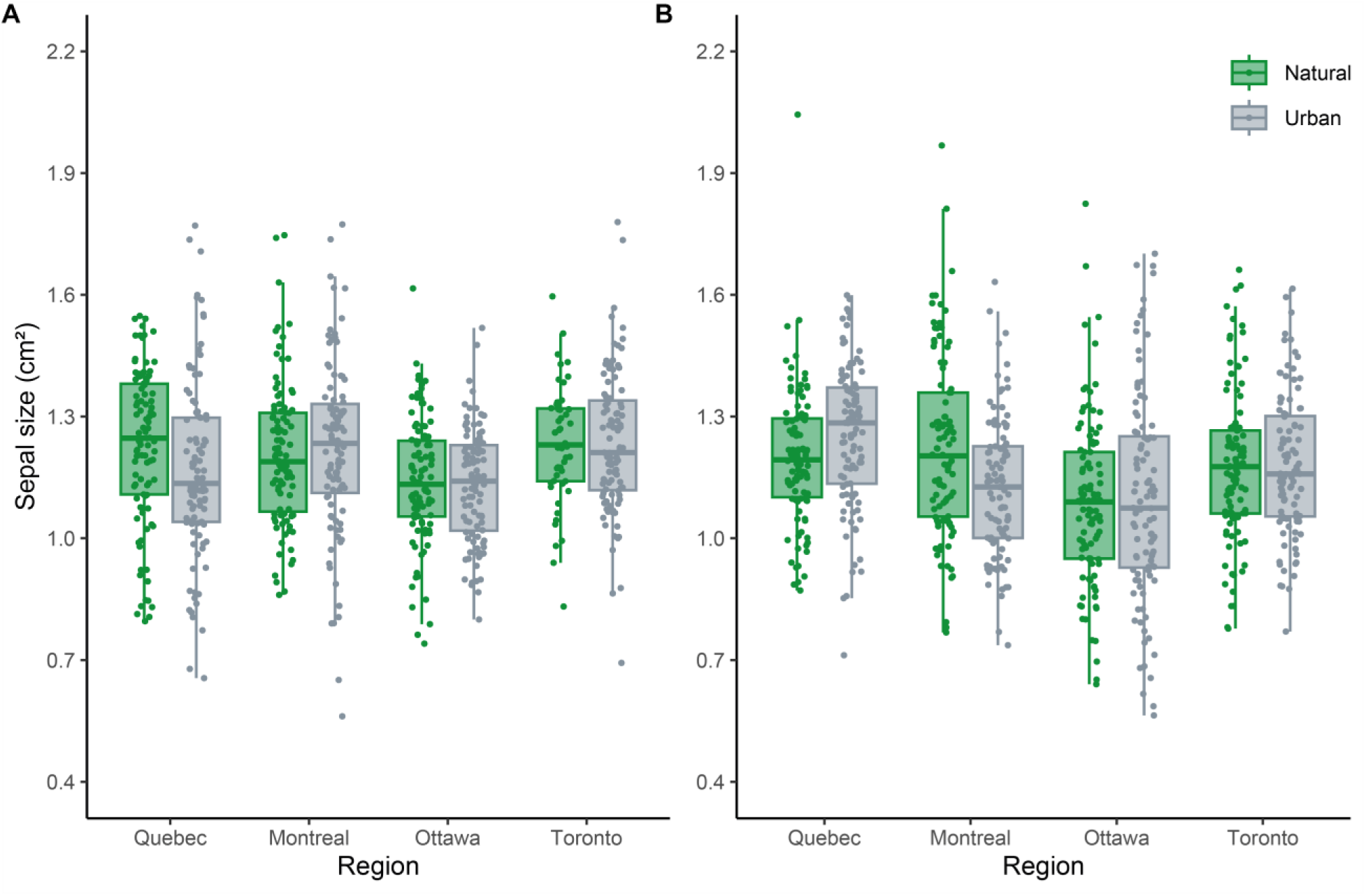
Sepal size in the field according to region and habitat type for 2021 (A) and 2022 (B). See Table S1 for more information about sampling.

**Fig. S4.**
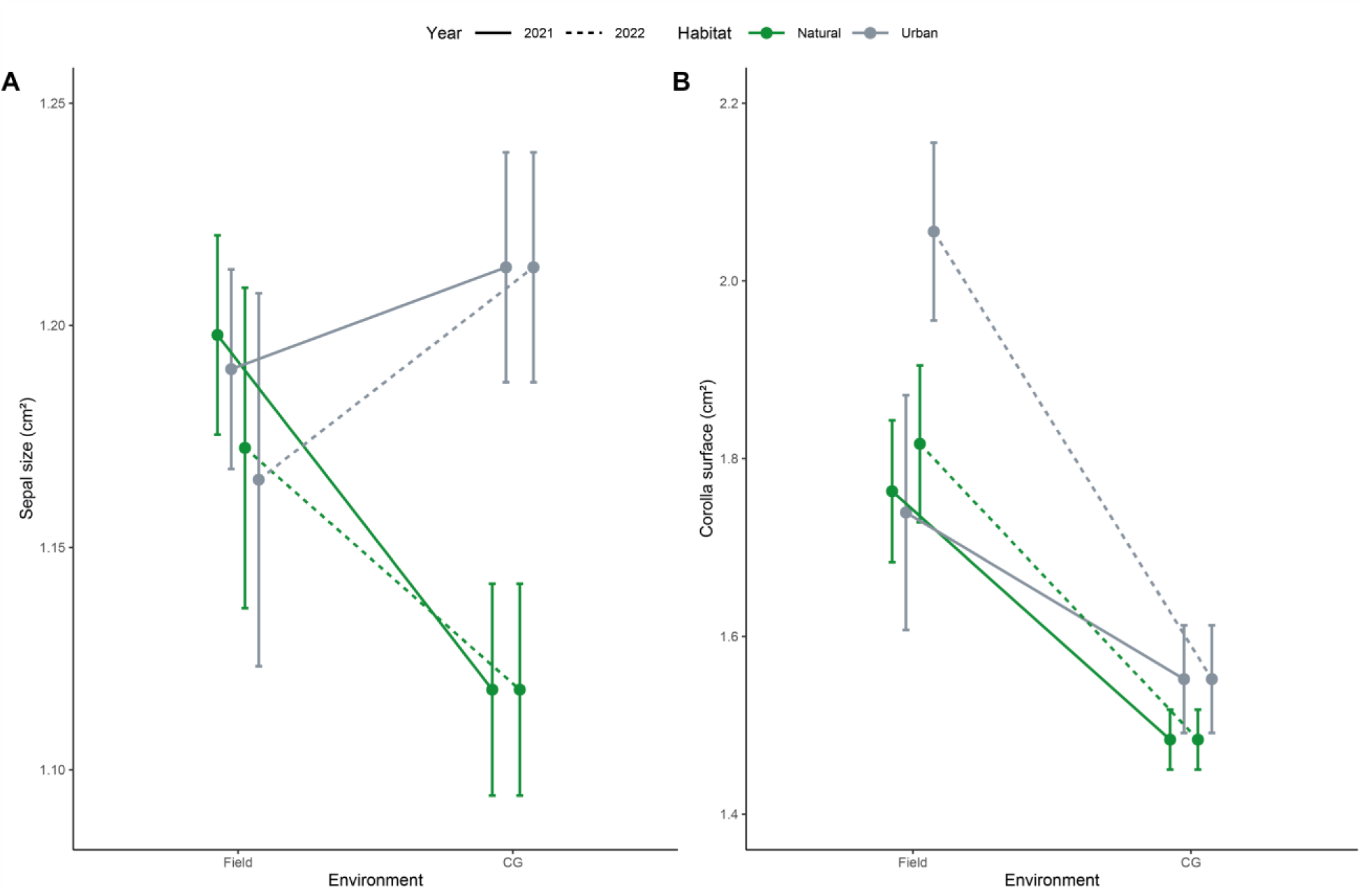
Mean response of sepal size (A) and corolla surface (B) to environment change, grouped by habitat. Solid lines represent 2021 data in the field compared to the unique common garden experiment, and dashed lines represent 2022 data in the field compared to the unique common garden experiment.

**Fig. S5.**
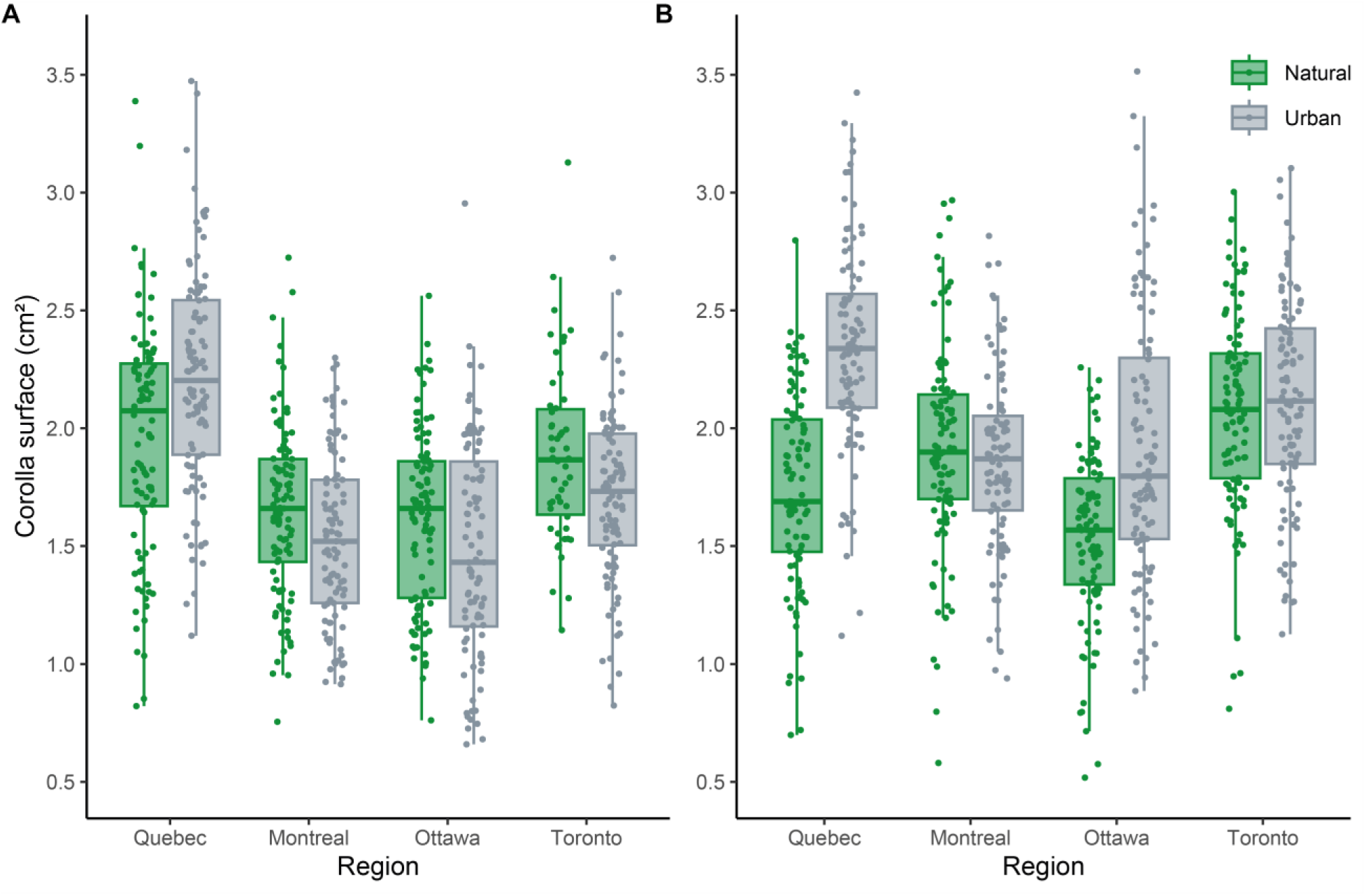
Corolla surface in the field according to region and habitat type for 2021 (A) and 2022 (B). See Table S1 for more information about sampling.

**Fig. S6.**
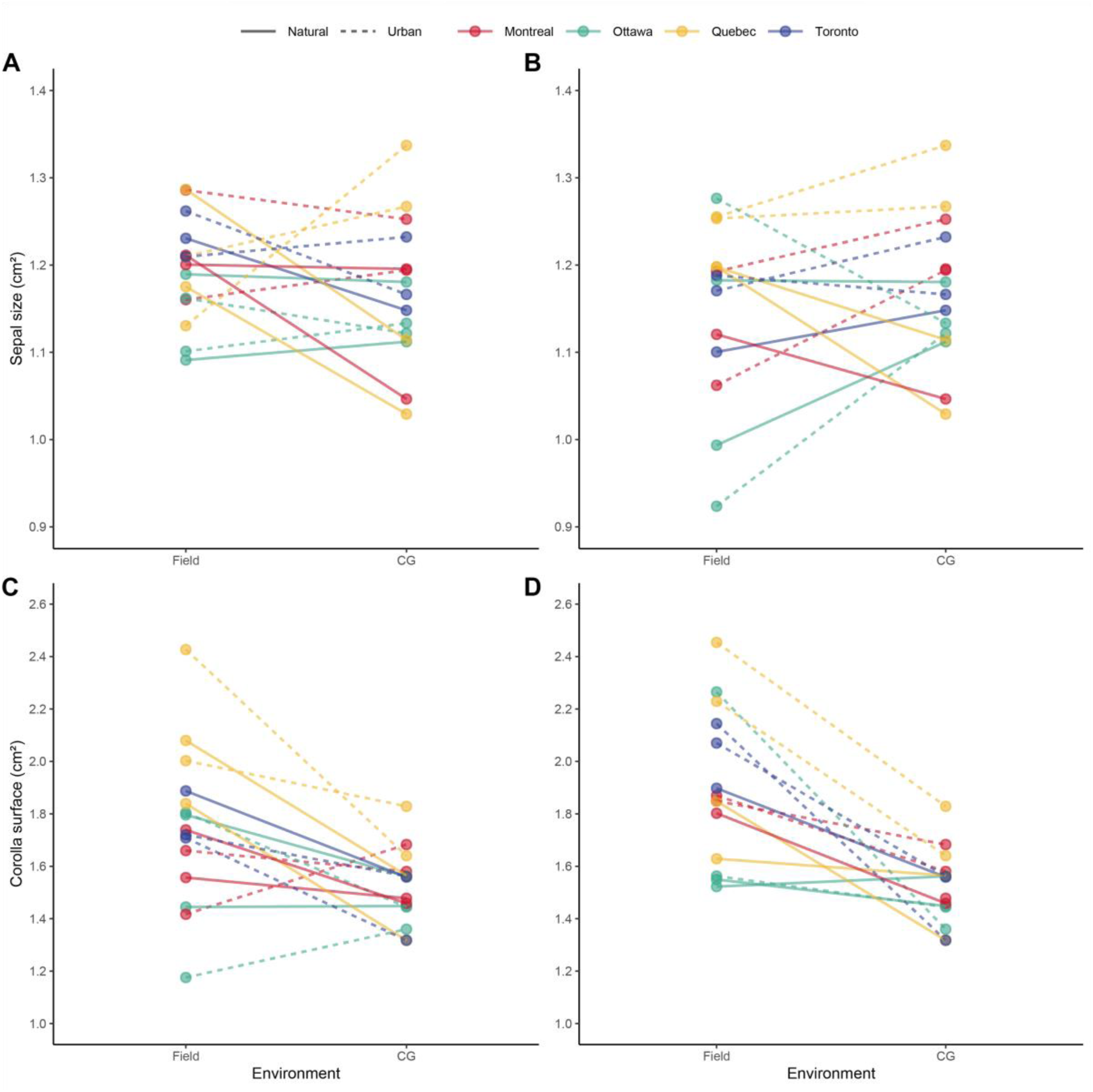
Plastic response of sepal size and corolla surface, grouped by region. Lines represent populations colored according to their region. Solid lines represent natural populations, and dashed lines represent urban populations. (A) Sepal size response for 2021 field data vs common garden data. (B) Sepal size response for 2022 field data vs common garden data. (C) Corolla surface response for 2021 field data vs common garden data. (D) Corolla surface response for 2022 field data vs common garden data.

**Fig S7.**
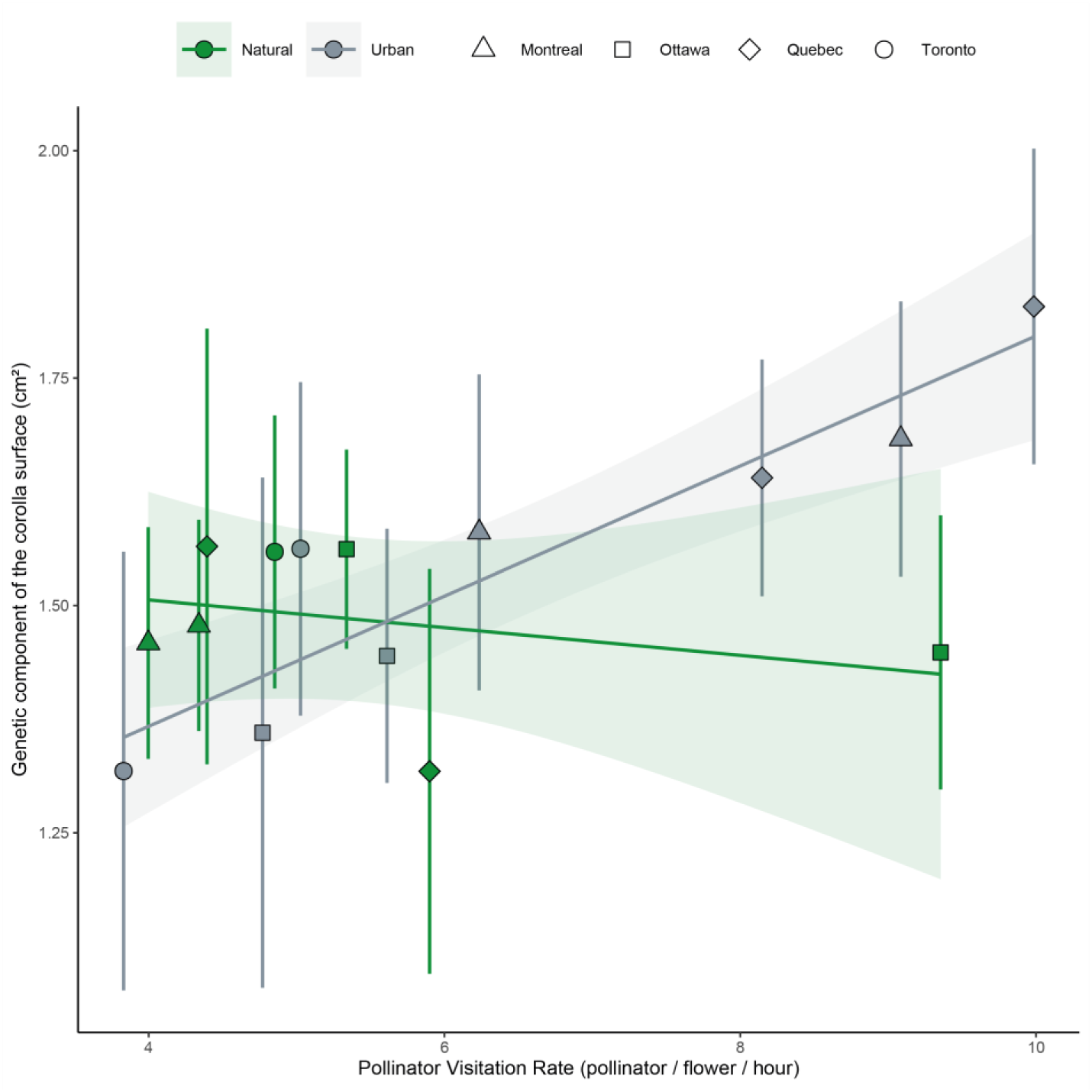
Correlation between the genetic component of the corolla surface and the pollinator visitation rate. There was a significant effect of the pollinator visitation rate on corolla surface, and a significant interaction between pollinator visitation rate and habitat. Regression lines for urban and natural populations are shown, with their standard errors. The points represent the population mean of the corolla surface in the common garden experiment (with 95% confidence intervals of the mean) and the population mean of the pollinator visitation rate averaged across the 2021 and 2022 field seasons.

## Supplementary Tables

**Table S1.**
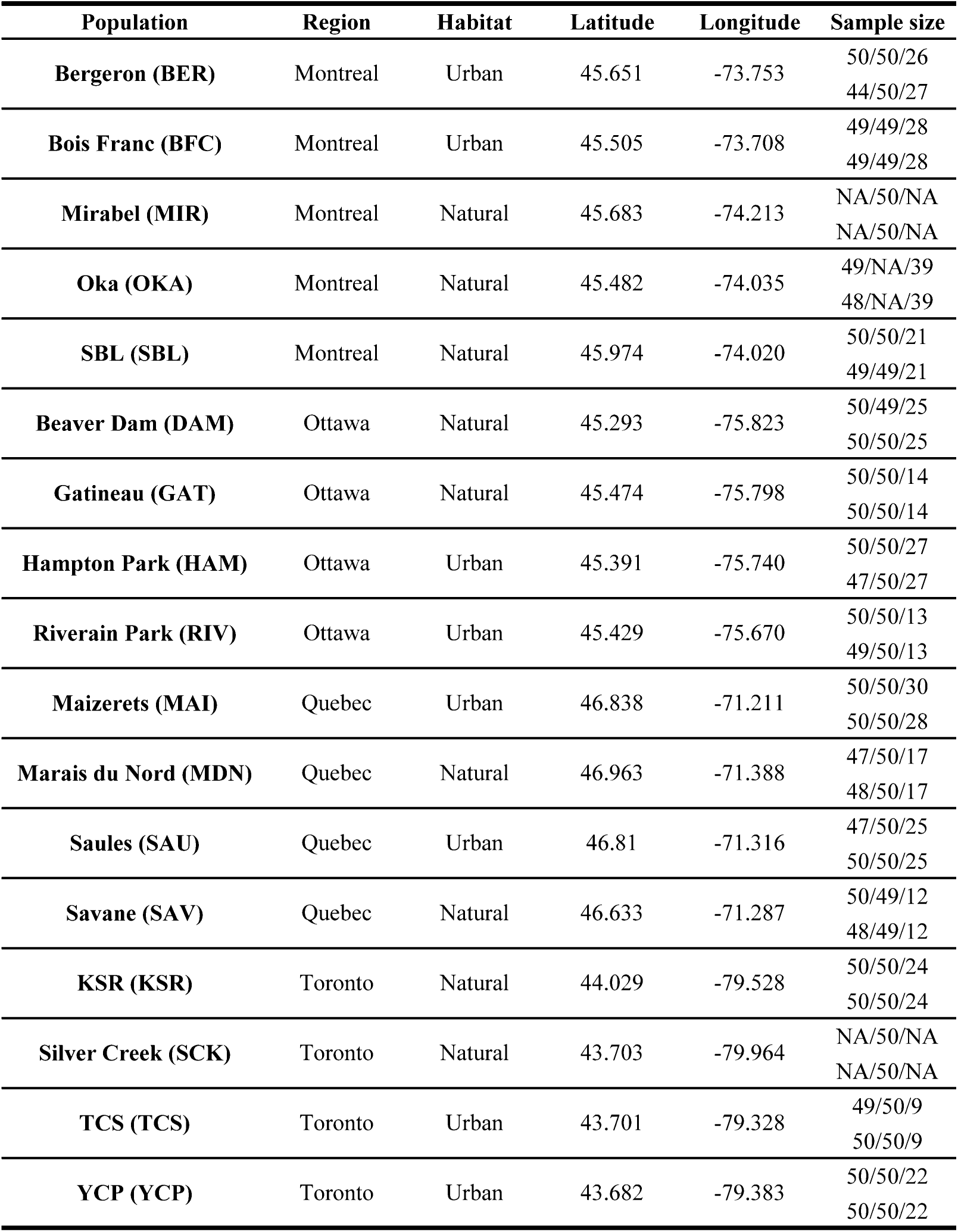
Populations studied. The first number in the sample size column is for the 2021 field season, the second one for the 2022 field season, and the third for the common garden experiment. The first line in the sample size column for each population is for profile pictures (sepal), and the second one is for front pictures (corolla).

| Population | Region | Habitat | Latitude | Longitude | Sample size |
| --- | --- | --- | --- | --- | --- |
| **Bergeron (BER)** | Montreal | Urban | 45.651 | -73.753 | 50/50/26<br>44/50/27 |
| **Bois Franc (BFC)** | Montreal | Urban | 45.505 | -73.708 | 49/49/28<br>49/49/28 |
| **Mirabel (MIR)** | Montreal | Natural | 45.683 | -74.213 | NA/50/NA<br>NA/50/NA |
| **Oka (OKA)** | Montreal | Natural | 45.482 | -74.035 | 49/NA/39<br>48/NA/39 |
| **SBL (SBL)** | Montreal | Natural | 45.974 | -74.020 | 50/50/21<br>49/49/21 |
| **Beaver Dam (DAM)** | Ottawa | Natural | 45.293 | -75.823 | 50/49/25<br>50/50/25 |
| **Gatineau (GAT)** | Ottawa | Natural | 45.474 | -75.798 | 50/50/14<br>50/50/14 |
| **Hampton Park (HAM)** | Ottawa | Urban | 45.391 | -75.740 | 50/50/27<br>47/50/27 |
| **Riverain Park (RIV)** | Ottawa | Urban | 45.429 | -75.670 | 50/50/13<br>49/50/13 |
| **Maizerets (MAI)** | Quebec | Urban | 46.838 | -71.211 | 50/50/30<br>50/50/28 |
| **Marais du Nord (MDN)** | Quebec | Natural | 46.963 | -71.388 | 47/50/17<br>48/50/17 |
| **Saules (SAU)** | Quebec | Urban | 46.81 | -71.316 | 47/50/25<br>50/50/25 |
| **Savane (SAV)** | Quebec | Natural | 46.633 | -71.287 | 50/49/12<br>48/49/12 |
| **KSR (KSR)** | Toronto | Natural | 44.029 | -79.528 | 50/50/24<br>50/50/24 |
| **Silver Creek (SCK)** | Toronto | Natural | 43.703 | -79.964 | NA/50/NA<br>NA/50/NA |
| **TCS (TCS)** | Toronto | Urban | 43.701 | -79.328 | 49/50/9<br>50/50/9 |
| **YCP (YCP)** | Toronto | Urban | 43.682 | -79.383 | 50/50/22<br>50/50/22 |

**Table S2.** Pollinator relative frequency calculated on collected specimens. The 2022 collection effort was more important (see Material and Methods), and this explains why more specimens were observed. NA are for species not found in either 2021 or 2022. Values are rounded.

| Species | Frequency |  |  |
| --- | --- | --- | --- |
|  | 2021 (n = 171 insects) | 2022 (n = 521 insects) | Cumulated (n=692 insects) |
| *Agapostemon virescens* | NA | 0.002 | 0.001 |
| *Ancistrocerus adiabatus* | NA | 0.002 | 0.001 |
| *Anthophora terminalis* | 0.070 | 0.036 | 0.045 |
| *Apis mellifera* | 0.088 | 0.056 | 0.064 |
| *Augochlora pura* | 0.018 | 0.015 | 0.016 |
| *Bombus borealis* | 0.012 | NA | 0.003 |
| *Bombus fervidus* | 0.006 | 0.002 | 0.003 |
| *Bombus impatiens* | 0.596 | 0.633 | 0.624 |
| *Bombus ternarius* | 0.018 | 0.002 | 0.006 |
| *Bombus vagans* | 0.099 | 0.136 | 0.127 |
| *Calliphoridae* | NA | 0.004 | 0.003 |
| *Ceratina calcarata* | NA | 0.002 | 0.001 |
| *Dolichovespula arenaria* | 0.023 | 0.012 | 0.014 |
| *Ectemnius continuus* | NA | 0.004 | 0.003 |
| *Epistrophe sp.* | NA | 0.002 | 0.001 |
| *Megachile mendica* | 0.006 | 0.002 | 0.003 |
| *Megachile relativa* | NA | 0.002 | 0.001 |
| *Muscidae* | NA | 0.002 | 0.001 |
| *Platycheirus nearcticus* | NA | 0.013 | 0.010 |
| *Rhingia nasica* | NA | 0.019 | 0.014 |
| *Toxomerus geminatus* | NA | 0.004 | 0.003 |
| *Unknown Syrphidae* | NA | 0.002 | 0.001 |
| *Unknown Vespula* | 0.006 | NA | 0.001 |
| *Vespula acadica* | 0.006 | NA | 0.001 |
| *Vespula alascensis* | NA | 0.002 | 0.001 |
| *Vespula consobrina* | 0.041 | 0.004 | 0.013 |
| *Vespula flavipilosa* | NA | 0.002 | 0.001 |
| *Vespula germanica* | NA | 0.004 | 0.003 |
| *Vespula maculifrons* | 0.012 | 0.036 | 0.030 |

**Table S3.**
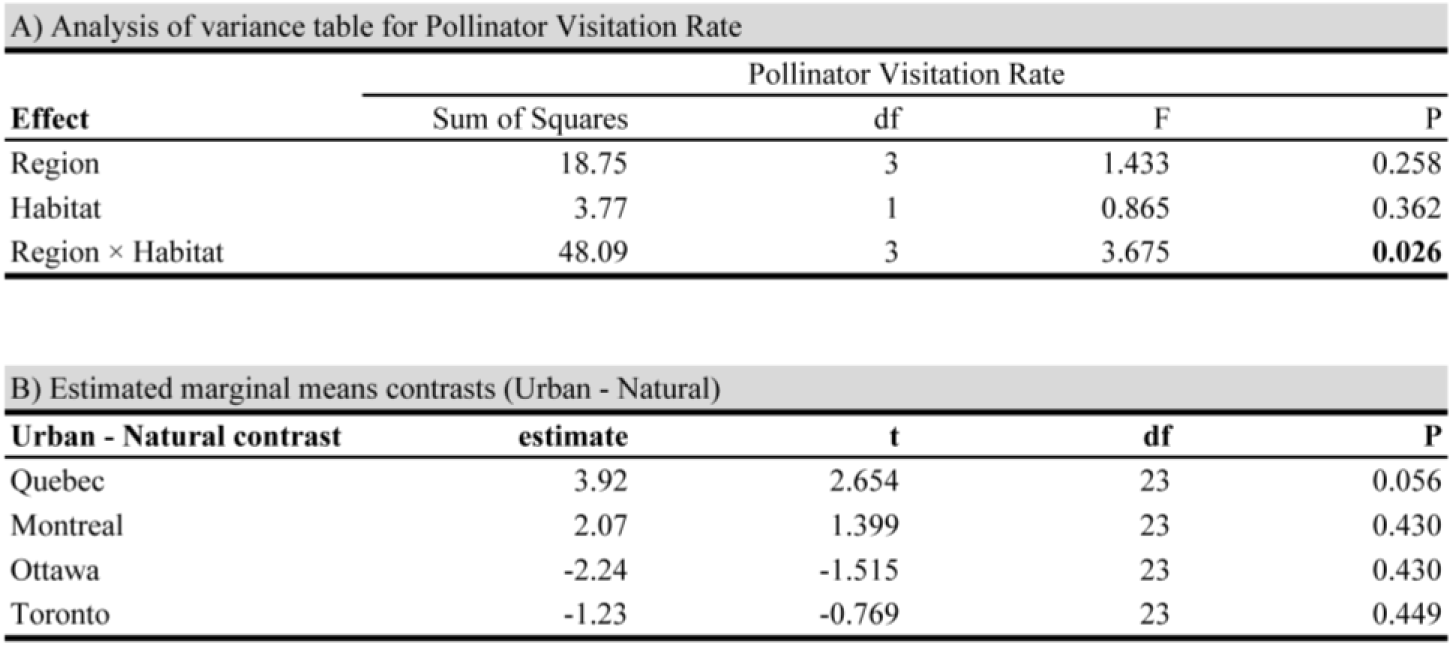
Analysis of the pollinator visitation rate variation.

**Table S4.**
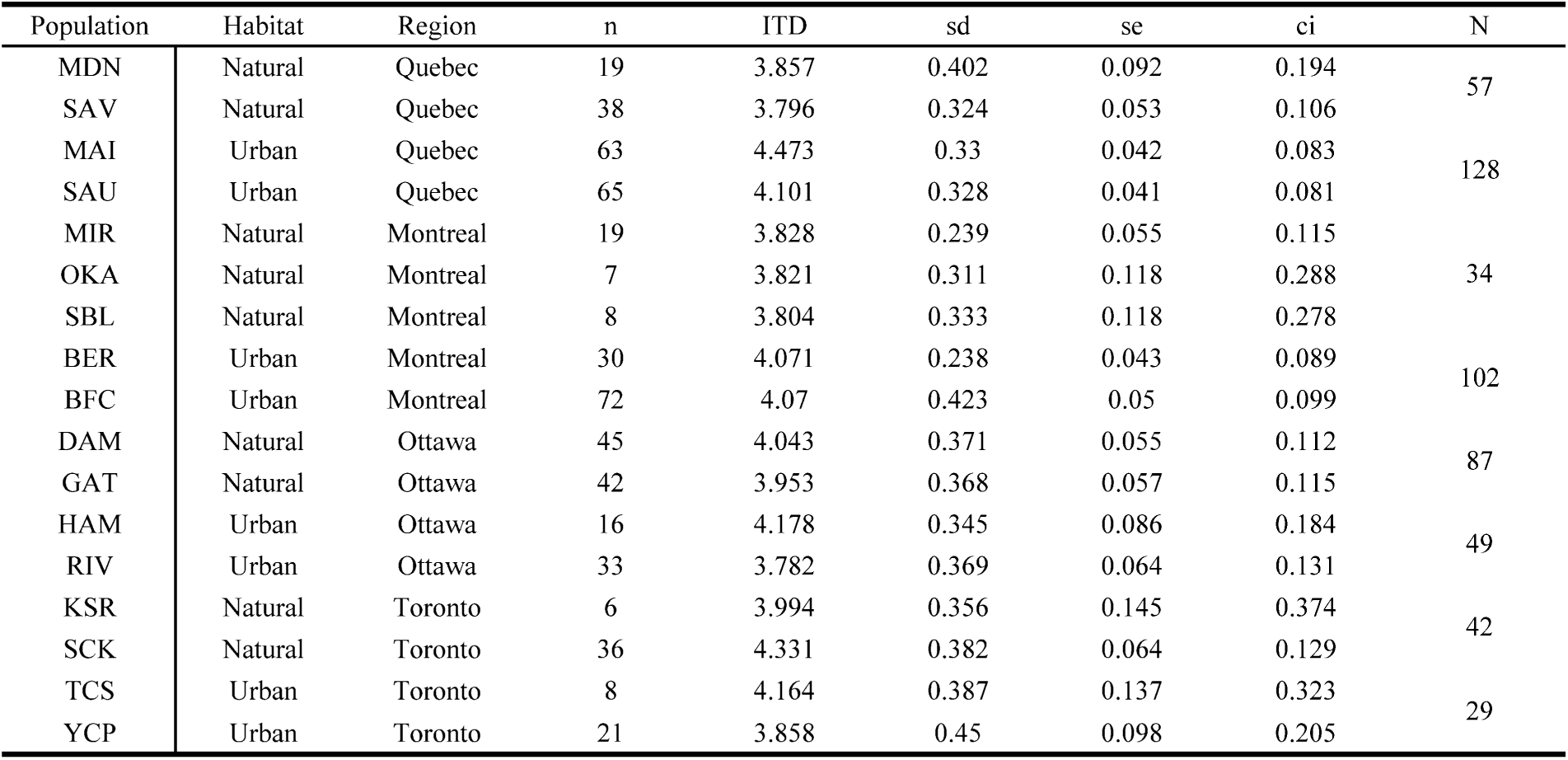
Sample size and summary of bumblebee sizes. . This table summarizes the 2021 and 2022 sampling years combined. n is the sample size for each population, and N is the sample size for each habitat within a region.

**Table S5.** Analysis of the bumblebee size variation.

| **A) Wald $\chi^2$ test** |  |  |  |
| --- | --- | --- | --- |
| **Effect** | Inter-Tegular Distance (ITD, mm) |  |  |
| | $\chi^2$ | df | P |
| Year | 0.109 | 1 | 0.74 |
| Region | 2.075 | 3 | 0.556 |
| Habitat | 2.414 | 1 | 0.12 |
| Year × Region | 1.989 | 3 | 0.574 |
| Year × Habitat | 0.302 | 1 | 0.582 |
| Region × Habitat | 8.826 | 3 | **0.031** |
| Year × Region × Habitat | 1.494 | 3 | 0.683 |

| **B) Estimated marginal means contrasts (Urban - Natural)** |  |  |  |  |
| --- | --- | --- | --- | --- |
| **Urban - Natural contrast** | Inter-Tegular Distance (ITD, mm) |  |  |  |
|  | estimate | t | df | P |
| Quebec | 0.45 | 2.599 | 7.21 | 0.138 |
| Montreal | 0.264 | 1.584 | 9 | 0.443 |
| Ottawa | -0.056 | -0.324 | 7.45 | 0.754 |
| Toronto | -0.279 | -1.141 | 28.29 | 0.526 |

We tested whether bumblebee inter-tegular distance (ITD) was explained by Year (2021 or 2022), Region (Quebec, Montreal, Ottawa, or Toronto), Habitat (Urban or Natural), and all interactions between these factors, using linear mixed models. **A)** Result of the Wald chi-square test **B)** We estimated the effect of habitat on inter-tegular distance for each region using post-hoc tests comparing estimated marginal means contrasts. We corrected for multiple comparisons using the holm correction.

**Table S6.** Analyses of the sepal size and corolla surface variation in the field.

| A) Wald $\chi^2$ test, male only | | | | | | |
| --- | --- | --- | --- | --- | --- | --- |
| Effect | Sepal Size |  |  | Corolla Surface |  |  |
| | $\chi^2$ | df | P | $\chi^2$ | df | P |
| Year | 8.385 | 1 | **0.003** | 33.597 | 1 | **<0.001** |
| Region | 3.834 | 3 | 0.279 | 37.237 | 3 | **<0.001** |
| Habitat | 0.344 | 1 | 0.557 | 2.191 | 1 | 0.138 |
| Year × Region | 28.021 | 3 | **<0.001** | 25.117 | 3 | **<0.001** |
| Year × Habitat | 8.146 | 1 | **0.004** | 54.977 | 1 | **<0.001** |
| Region × Habitat | 0.302 | 3 | 0.959 | 16.120 | 3 | **0.001** |
| Year × Region × Habitat | 6.980 | 3 | 0.072 | 10.690 | 3 | **0.013** |

| B) Estimated marginal means contrasts (Urban - Natural), male only |  |  |  |  |  |  |  |  |
| --- | --- | --- | --- | --- | --- | --- | --- | --- |
| Urban - Natural contrast | Sepal Size |  |  |  | Corolla surface |  |  |  |
|  | estimate | t | df | P | estimate | t | df | P |
| Quebec 2021 | -0.060 | -0.646 | 9.36 | 1.000 | 0.293 | 2.363 | 10.3 | 0.234 |
| Quebec 2022 | 0.067 | 0.719 | 9.43 | 1.000 | 0.623 | 4.992 | 10.6 | **0.004** |
| Montreal 2021 | -0.059 | -0.678 | 10.08 | 1.000 | -0.167 | -1.386 | 12.2 | 0.572 |
| Montreal 2022 | -0.068 | -0.781 | 10.15 | 1.000 | -0.007 | -0.056 | 12.4 | 1.000 |
| Ottawa 2021 | -0.027 | -0.291 | 9.45 | 1.000 | -0.223 | -1.776 | 10.8 | 0.416 |
| Ottawa 2022 | -0.008 | -0.080 | 9.48 | 1.000 | 0.364 | 2.905 | 10.7 | 0.103 |
| Toronto 2021 | -0.081 | -0.826 | 11.39 | 1.000 | -0.271 | -1.905 | 16.2 | 0.374 |
| Toronto 2022 | 0.012 | 0.130 | 9.66 | 1.000 | 0.018 | 0.140 | 11.4 | 1.000 |

| C) Wald $\chi^2$ test, male and female | | | | | | |
| --- | --- | --- | --- | --- | --- | --- |
| Effect | Sepal Size |  |  | Corolla Surface |  |  |
| | $\chi^2$ | df | P | $\chi^2$ | df | P |
| Year | 17.529 | 1 | **<0.001** | 45.646 | 1 | **<0.001** |
| Region | 3.720 | 3 | 0.293 | 33.065 | 3 | **<0.001** |
| Habitat | 0.239 | 1 | 0.624 | 2.279 | 1 | 0.131 |
| Year × Region | 24.444 | 3 | **<0.001** | 25.008 | 3 | **<0.001** |
| Year × Habitat | 6.151 | 1 | **0.013** | 63.618 | 1 | **<0.001** |
| Region × Habitat | 0.302 | 3 | 0.959 | 14.239 | 3 | **0.002** |
| Year × Region × Habitat | 6.806 | 3 | 0.078 | 6.638 | 3 | 0.084 |

| D) Estimated marginal means contrasts (Urban - Natural), male and female |  |  |  |  |  |  |  |  |
| --- | --- | --- | --- | --- | --- | --- | --- | --- |
| Urban - Natural contrast | Sepal Size |  |  |  | Corolla surface |  |  |  |
|  | estimate | t | df | P | estimate | t | df | P |
| Quebec 2021 | -0.060 | -0.672 | 9.48 | 1.000 | 0.255 | 2.005 | 10.4 | 0.358 |
| Quebec 2022 | 0.057 | 0.630 | 9.44 | 1.000 | 0.604 | 4.756 | 10.4 | **0.006** |
| Montreal 2021 | -0.044 | -0.527 | 10.14 | 1.000 | -0.183 | -1.501 | 12.1 | 0.636 |
| Montreal 2022 | -0.066 | -0.784 | 10.14 | 1.000 | 0.003 | 0.021 | 11.9 | 1.000 |
| Ottawa 2021 | -0.009 | -0.097 | 9.43 | 1.000 | -0.137 | -1.078 | 10.4 | 0.916 |
| Ottawa 2022 | 0.012 | 0.131 | 9.44 | 1.000 | 0.378 | 2.975 | 10.3 | 0.094 |
| Toronto 2021 | -0.068 | -0.728 | 10.92 | 1.000 | -0.311 | -2.234 | 14 | 0.254 |
| Toronto 2022 | -0.001 | -0.009 | 9.43 | 1.000 | 0.034 | 0.266 | 10.3 | 1.000 |

We tested whether sepal size or corolla surface was explained by Year (2021 or 2022), Region (Quebec, Montreal, Ottawa, or Toronto), Habitat (Urban or Natural), and all interactions between these factors using linear mixed models. Because the Sex factor was associated with sepal size and corolla surface in the field, we decided to run the same models with males only (**A-B**) and male and female (**C-D**) to assess the effect of females on our results. For each test (either male only or male and female combined), we estimated the effect of habitat on the sepal size and corolla surface for each region using post-hoc tests comparing estimated marginal means contrasts. We corrected for multiple comparisons using holm correction.

**Table S7.** Analyses of the sepal size and corolla surface variation in the common garden experiment.

| **A) Wald $\chi^2$ test** |  |  |  |  |  |  |
| --- | --- | --- | --- | --- | --- | --- |
| **Effect** | **Sepal size** |  |  | **Corolla surface** |  |  |
|  | **$\chi^2$** | **df** | **P** | **$\chi^2$** | **df** | **P** |
| Region | 2.270 | 3 | 0.084 | 5.113 | 3 | 0.163 |
| Habitat | 8.564 | 1 | **0.003** | 2.683 | 1 | 0.101 |
| Region × Habitat | 8.329 | 3 | **0.039** | 8.136 | 3 | **0.043** |

| B) Estimated marginal means contrasts (Urban - Natural) |  |  |  |  |  |  |  |  |
| --- | --- | --- | --- | --- | --- | --- | --- | --- |
| Urban - Natural contrast | Sepal Size |  |  |  | Corolla surface |  |  |  |
|  | estimate | t | df | P | estimate | t | df | P |
| Quebec | 0.227 | 3.700 | 7.29 | 0.028 | 0.281 | 2.617 | 7.78 | 0.126 |
| Montreal | 0.095 | 1.660 | 5.42 | 0.459 | 0.162 | 1.709 | 4.61 | 0.459 |
| Ottawa | -0.015 | -0.249 | 7.13 | 1 | -0.082 | -0.762 | 7.02 | 0.942 |
| Toronto | 0.043 | 0.585 | 6.91 | 1 | -0.081 | -0.623 | 6.84 | 0.942 |

We tested whether the sepal size or corolla surface was explained by Region (Quebec, Montreal, Ottawa, or Toronto), Habitat (Urban or Natural), and the interaction between these factors, using linear mixed models. **A)** Result of the Wald chi-square test **B)** We estimated the effect of habitat on the sepal size and corolla surface for each region using post-hoc tests comparing estimated marginal means contrasts. We corrected for multiple comparisons using holm correction.

**Table S8.** Analysis of the mean responses of sepal size and corolla surface, grouped by habitats, to environment change.

| Effect | Sepal Size |  |  |  |  |  | Corolla size |  |  |  |  |  |
| --- | --- | --- | --- | --- | --- | --- | --- | --- | --- | --- | --- | --- |
|  | 2021 |  |  | 2022 |  |  | 2021 |  |  | 2022 |  |  |
| | $\chi^2$ | df | P | $\chi^2$ | df | P | $\chi^2$ | df | P | $\chi^2$ | df | P |
| Environment | 0.916 | 1 | 0.338 | 0.047 | 1 | 0.837 | 8.859 | 1 | **0.002** | 36.761 | 1 | **<0.001** |
| Habitat | 2.003 | 1 | 0.156 | 4.052 | 1 | **0.044** | 1.445 | 1 | 0.229 | 2.492 | 1 | 0.114 |
| Environment × Habitat | 4.796 | 1 | **0.028** | 3.706 | 1 | 0.054 | 0.519 | 1 | 0.471 | 1.486 | 1 | 0.222 |

We tested whether the sepal size or corolla surface was explained by Environment (Field or Common Garden), Habitat (Urban or Natural), and the interaction between these factors, using linear mixed models. The comparison between the common garden and the 2021 field season represents the effect of genetic and plastic changes on the two components of flower size (in the common garden, we measure the offspring of seeds collected in the field in 2021). In 2022, we compared the same generation to one another (assessing the plasticity more than the genetic changes). Therefore, we ran separate models for 2021 and 2022.

**Table S9.**
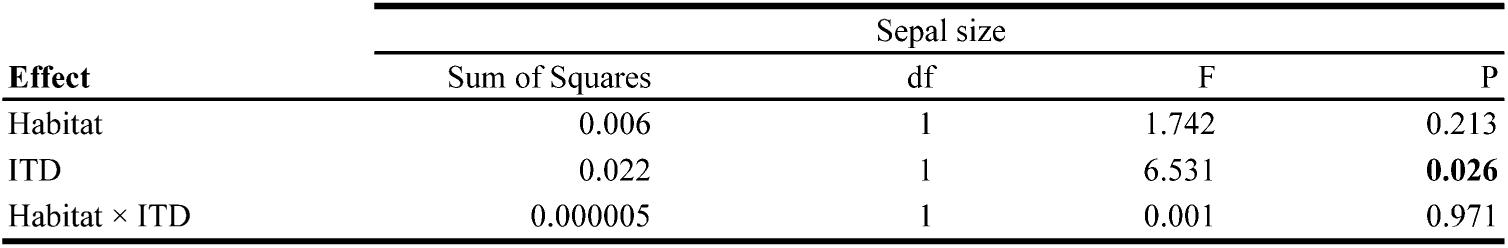
Analysis of the relationship between the genetic component of sepal size variation and bumblebee inter-tegular distances.

| Effect | Sepal size |  |  |  |
| --- | --- | --- | --- | --- |
|  | Sum of Squares | df | F | P |
| Habitat | 0.006 | 1 | 1.742 | 0.213 |
| ITD | 0.022 | 1 | 6.531 | **0.026** |
| Habitat × ITD | 0.000005 | 1 | 0.001 | 0.971 |

**Table S10.** Analysis of the relationship between the genetic component of corolla surface variation and the visitation rate of pollinators.

| **Effect** | Corolla surface |  |  |  |
| --- | --- | --- | --- | --- |
|  | Sum of Squares | df | F | P |
| Habitat | 0.001 | 1 | 0.288 | 0.601 |
| Pollinator Visitation Rate | 0.085 | 1 | 13.2 | **0.003** |
| Habitat × Pollinator Visitation Rate | 0.096 | 1 | 14.851 | **0.002** |

**Table S11.**
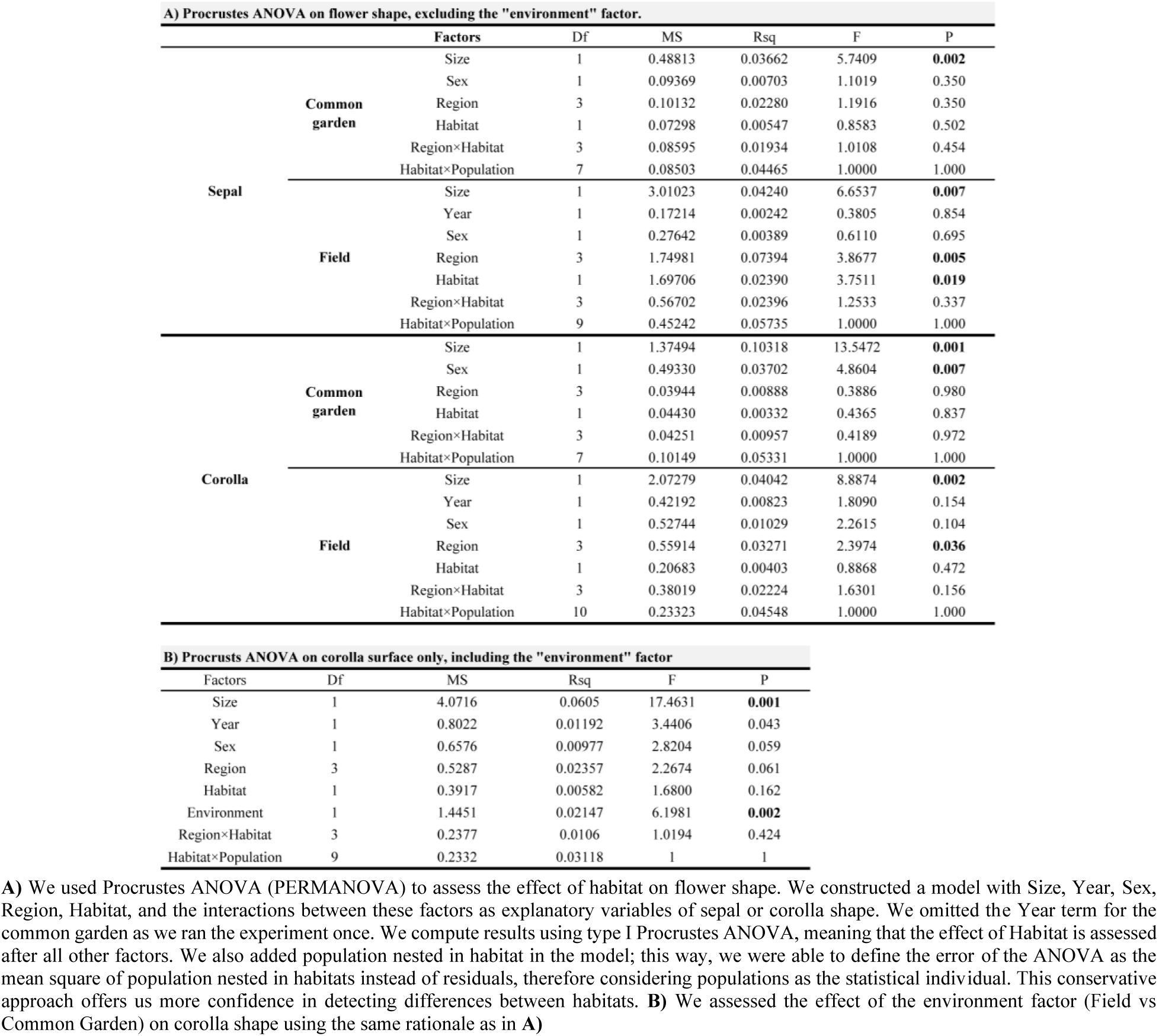
Analysis of the flower shape variation in the common garden experiment and in the field.

Tables S12-S22 are provided in supplementary file Table_S12_S22.xlsx

**Table S12. The effect of habitat, region, year and all interactions on bumblebee inter-tegular distance.** We show the estimate, standard error and t value for each factor level and interaction, compared to the intercept.

**Table S13. The effect of habitat, region and all interactions on sepal size in the common garden experiment.** We show the estimate, standard error and t value for each factor level and interaction, compared to the intercept.

**Table S14. The effect of habitat, region, year and their interaction on sepal size in the field, for male only flowers.** We show the estimate, standard error and t value for each factor level and interaction, compared to the intercept.

**Table S15. The effect of habitat, region, year and interactions on sepal size in the field, for male and female flowers.** We show the estimate, standard error and t value for each factor level and interaction, compared to the intercept.

**Table S16. The effect of environment, urbanization and their interaction on sepal size in 2021.** We show the estimate, standard error and t value for each factor level and interaction, compared to the intercept.

**Table S17. The effect of environment, urbanization and their interaction on sepal size in 2022.** We show the estimate, standard error and t value for each factor level and interaction, compared to the intercept.

**Table S18. The effect of habitat, region and all interactions on corolla surface in the common garden experiment.** We show the estimate, standard error and t value for each factor level and interaction, compared to the intercept.

**Table S19. The effect of habitat, region, year and their interaction on corolla surface in the field, for male only flowers.** We show the estimate, standard error and t value for each factor level and interaction, compared to the intercept.

**Table S20. The effect of habitat, region, year and interactions on corolla surface in the field, for male and female flowers.** We show the estimate, standard error and t value for each factor level and interaction, compared to the intercept.

**Table S21. The effect of environment, urbanization and their interaction on corolla surface in 2021.** We show the estimate, standard error and t value for each factor level and interaction, compared to the intercept.

**Table S22. The effect of environment, urbanization and their interaction on corolla surface in 2022.** We show the estimate, standard error and t value for each factor level and interaction, compared to the intercept.

